# Stress-Encoded Mitochondrial Plasticity: ATF4 Control of Mega-Mitochondria and Nanotunnel Communication

**DOI:** 10.64898/2026.03.04.709625

**Authors:** Amber Crabtree, Suraj Thapliyal, Mohd Mabood Khan, Edgar Garza Lopez, Andrea Marshall, Calixto Pablo Hernandez Perez, Oleg Kovtun, Jenny C. Schafer, Dea Pulatani, Yuho Kim, Sepiso K Masenga, Annet Kirabo, Jeremiah Afolabi, Melanie McReynolds, Renata O. Pereira, Prasanna Venkhatesh, Prasanna Katti, Brian Glancy, Antentor Hinton

**Affiliations:** Department of Molecular Physiology and Biophysics, Vanderbilt University, Nashville, Tennessee, USA; The Frist Center for Autism and Innovation, Vanderbilt University, Nashville, Tennessee, USA; Department of Physics and Astronomy, Vanderbilt University, Nashville, Tennessee, USA; Department of Biology, Indian Institute of Science Education and Research (IISER) Tirupati, AP, 517619, India; Department of Medicine, Division of Genetic Medicine & Clinical Pharmacology, Vanderbilt University Medical Center, Nashville, TN 37232, USA; Department of Internal Medicine, University of Iowa, Iowa City, IA, 52242, USA; Department of Biomedical Sciences, School of Graduate Studies, Meharry Medical College, Nashville, TN 37208-3501, USA; Department of Chemistry, Vanderbilt University, Nashville, TN, 37232, USA; Department of Cell and Developmental Biology, Vanderbilt University, Nashville, TN, 37232, USA; Department of Physical Therapy and Kinesiology, University of Massachusetts Lowell, Lowell, MA, USA; Department of Physiological Sciences, Pathology and Microbiology, HAND, research group, Mulungushi University School of Medicine and Health Sciences, Livingstone, Zambia; The Huck Institutes of the Life Sciences, Pennsylvania State University, University Park, PA, USA; Department of Biochemistry and Molecular Biology, Pennsylvania State University, University Park, PA, USA; Fraternal Order of Eagles Diabetes Research Center, Iowa City, IA, 52242, USA; National Heart, Lung, and Blood Institute, National Institutes of Health, Bethesda, MD; National Institute of Arthritis and Musculoskeletal and Skin Diseases, NIH, Bethesda, MD

**Keywords:** Stress, 3D Structure, Mitochondria, Metabolism, ATF4

## Abstract

Mitochondrial structural plasticity is a critical adaptive response to cellular stress, yet the transcriptional networks governing the formation of specialized mitochondrial architectures remain poorly defined. Here, we identified and demonstrated that activating transcription factor 4 (ATF4), the master regulator of the integrated stress response, directly regulates mitochondrial morphological remodeling through a novel ATF4-NRF1/Nrf2-MFN2 signaling axis. Using serial block-face scanning electron microscopy and three-dimensional reconstruction in *Drosophila* flight muscle, primary myotubes, and human skeletal muscle, we show that overexpression of ATF4 promotes significant mitochondrial elongation, increased cristae concentration, enhanced mitochondrial-endoplasmic reticulum contact site (MERC) formation, and the initiation of Mitochondrial Nanotunnels. In contrast, loss of ATF4 results in mitochondrial fragmentation and impaired aerobic capacity. Chromatin immunoprecipitation sequencing reveals direct ATF4 binding at the promoters of the genes encoding NRF1 and Nrf2, which in turn regulate *MFN2* expression. Small-molecule inhibition studies further establish that activation of this hierarchical pathway is both necessary and sufficient for stress-induced mitochondrial structural adaptation. Together, these findings position ATF4 as a master regulator of mitochondrial architectural plasticity, providing a direct mechanistic link between cellular stress signaling and organelle remodeling.

## Introduction

Mitochondria are essential organelles that produce cellular energy. However, recent advances in three-dimensional electron microscopy have revealed that these organelles possess substantially greater structural diversity and dynamic nature than the traditional “kidney bean” model. Mitochondrial structure plays an imperative role in determining function, and mitochondrial remodeling is evident in descriptions of distinct morphological phenotypes, which include donuts, Megamitochondria, nanotunnels, and highly branched networks (Vincent, Turnbull et al. 2017, Jenkins, Neikirk et al. 2024). These structural modifications are more than just physical oddities; they are becoming understood to have functional significance, allowing mitochondria to adapt to the cell’s changing metabolic, energy, and signaling needs.

Mitochondrial structure and function are highly connected. In addition to producing ATP, mitochondria regulate apoptosis, calcium homeostasis, redox signaling, and metabolic intermolecular coupling (Bratic and Larsson 2013, Del Campo, Contreras-Hernández et al. 2018). Mitochondria possess dynamic properties that include cycles of division mediated by proteins such as dynamin-related protein 1 (DRP1), as well as cycles of fusion mediated by proteins like optic atrophy protein 1 (OPA1) and mitofusin 1/2 (MFN1/2), which help accommodate the cell microenvironment (Chan 2012, Dong and Yao 2022). Any malfunction in these processes can lead to altered mitochondrial morphology and function, contributing to mitochondrial dysfunction in diverse pathological contexts (Bartsakoulia, Pyle et al. 2018)In addition to basal mitochondrial dynamics, mitochondria undergo substantial three-dimensional alterations in response to stress, adopting shapes that illustrate adaptive responses to proteostatic, metabolic, or oxidative stressors. (Glancy 2020).

There is mounting evidence that stress-induced mitochondrial morphologies are functionally specific. For example, although adaptive donut-shaped mitochondria may have a higher surface area to volume ratio, they may be less efficient at producing ATP and more suited for signaling and organelle interactions (Hara, Yuk et al. 2014, Cogliati, Enriquez et al. 2016). Expanded mitochondria-endoplasmic reticulum contact sites (MERCS) promote lipid metabolism, calcium exchange, and stress signaling (Bustos, Cruz et al. 2017). The formation of Megamitochondria is a stress-associated process shared by all organisms and tissues. On the other hand, Mitochondrial Nanotunnels enable long-range communication between mitochondria (Vincent, Turnbull et al. 2017, Hinton, Claypool et al. 2024, Jenkins, Neikirk et al. 2024). We still don’t know the regulatory logic governing the formation of these structures, conserved.

The integrated stress response (ISR) is a conserved signaling pathway that enables cells to adapt to diverse stressors, including endoplasmic reticulum stress, amino acid deprivation, hypoxia, and oxidative stress. The ISR activation leads to the phosphorylation of eIF2α through PERK, GCN2, PKR, or HRI, to trigger global depletion of protein synthesis and selective upregulation of ATF4 expression levels (Pakos-Zebrucka, Koryga et al. 2016). ATF4 is a multifunctional basic leucine zipper master transcriptional regulator, imparting a vital function in integrating the expression of genes involved in amino acid metabolism, antioxidant mechanisms, and autophagy in determining cell fate in a stress state context (Adams, Ebert et al. 2017, Rasmussen and Adams 2020). When activated, ATF4 heterodimerizes with diverse bZIP partners, allowing for combinatorial control.

Recent evidence suggests a possible association between the ISR and mitochondrial structural alterations. ER stress changes the architecture and function of mitochondria by affecting their lipid composition, metabolism, and MERCS (Hinton, Katti et al. 2024). PERK-stimulation of ATF4 has been shown to influence mitochondrial structural remodeling, indicating that ATF4 might function upstream of organelle structural adaptation (Jenkins, Neikirk et al. 2024). Given its critical role in coordinating metabolic and organelle stress responses, ATF4 has the potential to regulate long-term, transcriptionally encoded remodeling of mitochondrial structure. Here, we identify ATF4 as a master transcriptional regulator of mitochondrial architecture. Using three-dimensional electron microscopy, transcriptomics, and functional analyses across *Drosophila* and mammalian systems, we demonstrate that ATF4 orchestrates mitochondrial elongation, cristae densification, MERC expansion, and nanotunnel formation through a conserved NRF1/Nrf2–MFN2 axis (Vincent, Turnbull et al. 2017). We establish that ATF4 directly regulates the expression of nuclear respiratory factor 1 (NRF1), nuclear factor erythroid 2-related factor 2 (NRF2), and MFN, coordinating the formation of specialized mitochondrial structures during cellular stress. This study elucidates a mechanistic framework for understanding how cells employ specialized mitochondrial architectures to sustain cellular homeostasis under environmental stress, demonstrating the evolutionary conservation of this pathway from *Drosophila* to mammals.

## Materials and Methods

### Fly Model & Genetic Strains

The flies were maintained in vials or bottles on a standard yeast-cornmeal agar medium at 25 °C under a 12-hour light/dark cycle. The W1118 strain was used as the genetic background control for all the fly lines. The Mef2-Gal4 driver was used to achieve muscle-specific gene expression. The muscle-specific knockdown (KD) experiment used Mef2-Gal4 (Bloomington Stock #27390) flies crossed with Tub-Gal80ts at 18°C, which was later raised to 29°C after the larval stage was reached. When manipulating ATF4 signaling, RNAi knockdown and overexpression lines targeting ATF4/cryptocephal (crc) were crossed with Mef2-Gal4. Visualization of mitochondria was conducted using UAS-mito-GFP. All Drosophila stocks were procured from the Bloomington Drosophila Stock Center, and chromosomal designations and gene nomenclature followed FlyBase conventions (http://flybase.org).

### Mitochondrial Staining and Confocal Imaging

The thoraces of *Drosophila*, aged between 2-3 days were dissected in 4% paraformaldehyde (PF; Sigma-Aldrich) and indirect flight muscles were isolated as previously stated (Katti, Rai et al. 2021). The isolated muscles were then fixed 4% PF for 1.5 hours under constant stimulation, followed by three 15-minute washes in PBSTx (PBS containing 0.3% Triton X-100). We also used GFP expressed from UAS-mito-GFP under Mef2 Gal4 control. The experimenter labeled F actin by incubating muscles with 2.5 µg/mL phalloidin TRITC (Sigma Aldrich) in PBS for 40 minutes at room temperature. After staining, we mounted the muscles with Prolong Glass Antifade Mountant, with NucBlue (ThermoFisher). Then, we imaged the muscles on a Zeiss LSM 780 microscope.

For the distribution analysis, inside myotubes, we stained the cells with MitoTracker Orange to visualize mitochondrial morphology and subcellular localization. We also used methyl ester (TMRM) to measure the membrane potential to assess mitochondrial function. Spinning-disk super-resolution by Optical Pixel Reassignment (SoRa) microscopy was used to obtain super-resolution images. The intensity of radial fluorescence profiles was quantified across five concentric subcellular zones: nuclear, perinuclear, central, radial, and distal.

To obtain high-resolution z-stacks of live cells stained with MitoTracker Orange or MitoTracker Green/TMRM, confocal images were acquired using a 100× Plan Apo NA 1.45 objective on a spinning disk confocal system based on a Nikon Ti2-E inverted fluorescence microscope equipped with a Yokogawa CSU-W1, Hamamatsu Fusion BT camera, SoRa super-resolution module, environmental chamber, piezo stage controller, and solid-state laser diodes (405, 488, 561, and 640 nm), all controlled using NIS-Elements software (version 5.42; Nikon, Melville, NY). Emission filters were 605/52 nm for the 561 nm channel and 525/36 nm for the 488 nm channel. Serial optical sections were acquired in W1 confocal mode with a 300 nm z-step size.

### *Drosophila* Sample Transmission Electron Microscopy

The dissected samples comprised 2–3-day-old flies fixed with a common fixative solution of 2.5% glutaraldehyde, 1% paraformaldehyde, and 0.12 M sodium cacodylate buffer. Each flight muscle was then isolated and transferred to fresh fixative before processing. Mitochondrial and cristae morphology were manually traced and analyzed using the freehand tool NIH ImageJ software (Schneider, Rasband et al. 2012). Multi-Measure ROI tool in ImageJ was used to make measurements of mitochondrial area, circularity, length, and number (Lam, Katti et al. 2021, Neikirk, Lopez et al. 2023). At identical magnification three specific regions were examined to determine cristae area, circularity index, number, and cristae score. Estimation of cristae volume was conducted by dividing the total cristae area by the total mitochondrial area (Patra, Kwon et al. 2016). Mitochondria-endoplasmic reticulum contact sites (MERCS) were quantified by measuring MERC distance, MERC length, and percentage.

### Scanning Transmission Electron Microscope

Sections of epoxy-embedded tissues, measuring 60-80 nm in thickness, were excised using a diamond knife attached to an ultramicrotome (Leica UC7) and then placed on copper grids covered with formvar. To with lead citrate. A scanning transmission electron microscope (STEM) running at 200 kV was used for the imaging. Mitochondrial ultrastructure was visualized using both bright-field (BF) and high-angle annular dark-field (HAADF) detectors. Z-contrast, provided by HAADF imaging, further enhanced visibility. Images were captured at magnifications ranging from 20,000× to 150,000×, with pixel sizes of 0.5-2 nm and dwell times per pixel of 1-5 μs to reduce beam-induced damage.

Mitochondrial morphological metrics, such as length, MERC, were measured using ImageJ for quantitative analysis. The organization of the mitochondrial network was evaluated by measuring its length and branching, and the membrane proximity (<30 nm) was used to identify the mitochondria-ER contact points. With a minimum of three independent samples per condition and ten to twenty mitochondria evaluated per cell, all studies were conducted in a blinded fashion across three or more biological replicates.

### Animal Models

We used male C57BL/6J mice (n = 5-7) as animal models in each age group. We produced Atf4^fl/fl^ mice using the procedures that were previously described (Ebert, Dyle et al. 2012). Dr. Pierre Chambon of the University of Strasbourg generously donated the tamoxifen-inducible HSA-CreERT2 mice. The Atf4^fl/fl^ mice were generated by crossing Atf4 with HSA-CreERT2 mice. The mice were given free access to food and water and kept in a typical environment with a 12-hour light-dark cycle. The mice were euthanized using cervical dislocation followed by CO_2_ anesthesia. All serum samples were collected between 8:00 am and 12:00 pm. Total body weight was measured at the time of tissue collection. All animal-related procedures were conducted in compliance with the standards set forth by the institutional animal care and use committee.

### Primary Cell Culture

Satellite cells were isolated following previously verified methods (Pereira, Tadinada et al. 2017). Culture dishes coated with BD Matrigel were used to plate primary satellite cells that were extracted from Atf4^fl/fl^ mice (n = 8 for eachisolated from Atf4fl/fl mice (n = 8 per experimental condition). Isolated cells were then plated on BD Matrigel-coated dishes and cultured in DMEM/F-12 medium (Gibco, Waltham, MA). The plates were subsequently supplemented with 20% fetal bovine serum (FBS; Gibco), 40 ng/mL basic fibroblast growth factor, 1× non-essential amino acids, 0.14 mM β-mercaptoethanol, 1% penicillin/streptomycin, and 300 μL/100 mL Fungizone. After plating was complete, the growth factor concentration for basic fibroblast growth factor was reduced to 10 ng/mL to maintain it. When 90% confluency was reached, differentiation was activated by switching DMEM/F-12 containing 2% FBS and 1× insulin-transferrin-selenium.

Three days after differentiation, myotubes were infected with an adenovirus expressing GFP-Cre to generate ATF4 knockout (ATF4 KO) myotubes. An ATF4-expressing adenovirus (Ad5CMV-ATF4/RSVeGFP) was used to infect differentiated myotubes to produce ATF4 overexpression (ATF4 OE) for gain-of-function studies. Measurements for all experiments were performed at 3 and 7 days post-viral infection. The culture medium was replaced on alternating days by washing with phosphate-buffered saline (PBS) to remove residual media. Maintenance of C2C12 myoblasts was carried out at 37°C in a humid atmosphere containing 5% CO2. The growth medium consisted of DMEM containing 4.5 g/L glucose supplemented with 10% fetal bovine serum. The medium was replaced with DMEM containing 2% horse serum after the cells reached confluency to induce differentiation into myotubes. After five days of differentiation initiation, the myotubes were treated with either ethanol, 0.1 μM oligomycin, or 1 μg/mL tunicamycin ± 500 μM 4-phenylbutyric acid (PBA) for 8 hours.

### Murine and Human Sample Transmission Electron Microscopy

Cells were fixed in 2.5% glutaraldehyde in sodium cacodylate buffer for a period of 1 hour at 37°C as per the previous protocols (Hinton, Katti et al. 2023). Fixed samples were further embedded in 2% agarose, post-fixed in 1% osmium tetroxide in buffer, and stained with 2% uranyl acetate. using a graded series of ethanol. Resin embedding was done using EMbed 812. Sections of 80 nm thickness were prepared by an ultramicrotome and stained with 2% uranyl acetate and lead citrate. Microscopy was carried out on a JEOL JEM-1230 transmission electron microscope at an acceleration voltage of 120 kV.

### Serial Block Face-Scanning Electron Microscopy

We have prepared the samples according to previously published protocol (Garza-Lopez, Vue et al. 2021, Vue, Neikirk et al. 2023). The tissue samples were fixed in a 2% glutaraldehyde solution dissolved in 0.1 M cacodylate buffer before heavy metal staining, as previously published (Mustafi, Kikano et al. 2014). The samples underwent a sequential process that started with a 3% potassium ferrocyanide solution containing 2% osmium tetroxide at 4°C for 1 hour. The samples were filtered after the thiocarbohydrazide solution at 0.1% concentration was applied for 20 minutes. The samples were treated for 30 minutes with a 2% osmium tetroxide solution, then placed in 1% uranyl acetate solution for overnight storage at 4°C. The process required deionized water washes to be performed between each step. The following day, samples were treated with 0.6% lead aspartate solution (30 minutes at 60°C) and dehydrated through graded acetone dilutions. Tissues were infiltrated with TAAB 812 hard epoxy resin (Aldermaston, Berks, UK), embedded in fresh resin, and polymerized at 60°C for 36-48 hours. Following polymerization, blocks were sectioned for transmission electron microscopy to identify regions of interest, then trimmed to 0.5 mm × 0.5 mm and mounted on aluminum pins. Pins were placed in a scanning electron microscope for serial block-face imaging. Images were collected from 300-400 thin (0.09 μm) serial sections for three-dimensional reconstruction.

### 3D Reconstruction and Segmentation

Three-dimensional reconstruction of serial block-face scanning electron microscopy data was performed according to conventional protocols (Crabtree et al., 2023; Garza-Lopez et al., 2022; Hinton et al., 2023; Neikirk et al., 2023; Vue et al., 2023). A volume of data, approximately 300-400 orthoslices thick, comprising 50-100 sections was assembled for reconstruction in Amira software (ThermoFisher Scientific). An investigator blinded to experimental conditions and experienced organelle identification manually segmented structural features by tracing contours through sequential slices of micrograph blocks using contour-tracing tools. The resulting volumetric structures were quantified and assembled into three-dimensional visualizations (Garza-Lopez, Vue et al. 2021). For *Drosophila* samples, Imaris image analysis software was additionally employed for 3D reconstruction and volumetric quantification of mitochondrial networks.

### Western Blotting

Protein extracts isolated from differentiated myotubes as well as C2C12 cells, were obtained following approved protocols (Hinton, Katti et al. 2024). Cold PBS was used to wash cells, which were subsequently lysed in a cold lysis solution containing 25 mM Tris-HCl pH 7.9, 5 mM MgCl2, 10% glycerol, 100 mM KCl, 1% NP-40, 0.3 mM dithiothreitol, 5 mM sodium pyrophosphate, 1 mM sodium orthovanadate, 50 mM sodium fluoride, and a protease inhibitor cocktail. After lysis, cells were scraped from culture dishes, homogenized with a 25-gauge needle, and centrifuged at 14,000 rpm (5,268 × g) for 10 minutes at 4°C. Supernatants, were then diluted with Laemmli sample buffer to 1× concentration.

Protein was isolated by sodium dodecyl sulfate-polyacrylamide gel electrophoresis and transferred to nitrocellulose membranes (Bio-Rad, Berkeley, CA). Membranes were blocked using a solution of 5% bovine serum albumin in Tris-buffered saline containing 0.05% Tween-20. The primary antibodies that were used were BIP (BD Biosciences, #610978, 1:5,000), GRP75 (Cell Signaling Technology, #D13H4, 1:1,000), VDAC (Cell Signaling Technology, #4866, 1:1,000), ATF4 (Proteintech, Rosemont, IL, 10835-1-AP, 1:1,000), IP3R3 (BD Biosciences, #610312, 1:1,000), FGF21, OPA1 (BD Biosciences, San Jose, CA, #612606, 1:1,000), DRP1, MFN-1 (Abcam, Cambridge, UK, ab57602, 1:1,000), and MFN-2 (Abcam, ab101055, 1:1,000). GAPDH (Cell Signaling Technology, Danvers, MA, #2118, 1:5,000) and actin served as loading controls. For the secondary antibodies Alexa Fluor anti-rabbit 680 (Invitrogen, #A27042, 1:10,000) was included. The respective protein bands observed were quantified using Image Studio Lite Ver 5.2 software.

### RNA Extraction and Quantitative Real-Time PCR

RNA was extracted from the cells by employing the RNeasy Kit, as recommended by the supplier (Qiagen Inc.). Concentration and purity of RNA were determined by analyzing A260/A280 values by employing the NanoDrop 1000 spectrophotometer (NanoDrop Products, Wilmington, DE). Reverse transcriptions of 1 μg isolated RNA were performed by employing the High-Capacity cDNA Reverse Transcription Kit (Applied Biosystems, Carlsbad, CA), as earlier described (Boudina, Sena et al. 2007).

Real-time q-PCR was conducted employing SYBR Green dye (Life Technologies Inc.; Carlsbad, CA). The samples in each experimental group were run in triplicates with approximately 50 ng of cDNA per reaction, in a 384-well plate format, employing ABI Prism 7900 HT sequencing detection systems (Applied Biosystems). The cycling thermal profiles began with a denaturation step of 10 minutes at 95°C, followed by 40 cycles of amplification, each consisting of 15 seconds at 95°C, 15 seconds at 59°C, and finally an extension of 30 seconds at 72°C. Gene expression was determined by ΔΔCt analysis, normalized to GAPDH expression.

For *Drosophila* samples, RNA was isolated from the flight muscles of 1 2 day old flies. We removed the flight muscles from cut thoraces at 4 °C. Placed the flight muscles, in TRIzol reagent (Sigma Aldrich/Invitrogen). We extracted the RNA with a tissue grinder. Reverse transcription of the RNA was performed using SuperScript III (Invitrogen). For RT PCR we used 50 ng of cDNA per reaction. For RT PCR we mixed the cDNA with iTaq Universal SYBR Green (Bio Rad). Ran the reaction, on a QuantStudio 6 Flex system (Applied Biosystems).

### RNA Sequencing and Bioinformatic Analysis

Sequencing of RNA was done following previously established protocols (Hall, Medina et al. 2017). For *Drosophila* studies, RNA was isolated from five wild-type and five ATF4 knockdown flies. The sequencing reads were uploaded to Kallisto and quantified using Kallisto (v0.46.2) with an index built from BDGP6.28 genome assembly transcript sequences. Differential gene expression analysis was conducted using DESeq2 (v1.30.0). Gene list that exhibited expression levels greater than 5 fragments per kilobase per million reads were compiled and classed in either control or ATF4 knockdown samples.

For the mammalian cell line analysis, genes with differential expressions were found with the filter criterion of the absolute log2 fold change > 0.66 and an adjusted p-value < 0.05. For enrichment analyses, the tool used was Ingenuity Pathway Analysis (IPA) and WebGestalt for Gene Set Enrichment Analysis (GSEA). For IPA and GSEA, significance was assessed using the following: IPA: an adjusted p value of < 0.05. GSEA: an absolute Z-score of > 2 with an adjusted p value of < 0.05, and an absolute enrichment score of > 2. The PANTHER database (http://pantherdb.org/) was utilized to evaluate Gene Ontology data, whereas WebGestalt (http://www.webgestalt.org/) was used for pathway analysis, looking for KEGG pathways, and searching the gene promoters for binding sites of every transcription factor.

### Chromatin Immunoprecipitation Sequencing

ChIP-sequencing was performed following a 10-hour treatment with tunicamycin (2 μg/mL) in Atf4^+/+^, ATF4^-/-^, Chop^+/+^, and Chop^-/-^ MEFs, as described in previous literature (Han, Back et al. 2013, Tameire, Verginadis et al. 2019, Zou, Ohta et al. 2022). The DNA fragments obtained were processed for sequencing, which was performed according to the protocol. The peaks for ATF4 and CHOP binding at gene promoters were identified using standard peak identification algorithms.

### Mitochondrial Respiratory Function

Cell metabolism measurements were performed by using an XF24 Extracellular Flux Analyzer (Agilent Technologies-Seahorse Bioscience, North Billerica, MA), seeding a final concentration of 20,000 cells/well and differentiation as described above. On day three post-differentiation, cells were infected with adenoviruses expressing ntGFP and/or GFP-Cre at a high MOI calculated to achieve infection efficiencies higher than 95% as determined by fluorescence intensity. On day three post-infection, XF culture medium replaced regular culture medium, and the cells were placed in a non-CO_2_ incubator for 60 minutes.

The rate of basal oxygen consumption rate (OCR) was assessed using XF-DMEM, after which additional assessments of OCR were conducted in response to the following compounds added into the medium: oligomycin (1 μg/mL) to estimate ATP-linked respiration, FCCP (1 μM) to estimate maximum respiratory capacity, and a combination of rotenone (1 μM) and antimycin A (10 μM) to estimate non-mitochondrial respiration (Wende, O’Neill et al. 2015). The spare respiratory capacity was then calculated by subtracting basal OCR from maximum OCR.

### Measurement of Mitochondria Membrane Potential and Reactive Oxygen Species

The mitochondrial membrane potential was assessed using the fluorescent cationic dye tetramethylrhodamine methyl ester (TMRM). The quantification of TMRM fluorescence intensity loss over time within a given microscopic field was performed. The production of mitochondrial hydrogen peroxide was measured using MitoPeroxy Yellow 1 (MitoPY1). The TMRM fluorescence was quantified using the FIJI/ImageJ (NIH) method. The time-lapse image stacks were initially imported and, if needed, drift was corrected using the StackReg/TurboReg plugin. Regions of interest (ROIs) belonging to particular cells or mitochondrial-rich areas were manually identified and the mean fluorescence intensity for each ROI was calculated at all time points. All data were adjusted for background fluorescence, which was calculated from areas devoid of cells. To get the relative changes in mitochondrial membrane potential over time (F/F), the intensity values of the fluorescence were normalized to the original baseline fluorescence (F). Data from several ROIs and separate experiments were averaged for population analysis, and the fluorescence decay rate was determined to quantify mitochondrial depolarization kinetics. The data are represented as mean ± standard error of the mean from at least three separate experiments for each experimental group, with four fields per well at various time points.

### Calcium Imaging

Intracellular calcium dynamics were assessed in primary mouse myotubes using Fura-2 AM, a ratiometric imaging technique that alternates excitation at 340 nm and 380 nm, with emission at 510 nm. Calcium transients were induced and recorded over a time period of 300 seconds. Peak amplitude and area under the curve were quantified to assess calcium handling capacity across experimental groups.

### Statistical Analysis

Results are expressed as mean values ±SE of mean. If two groups were compared, unpaired Student’s *t* tests were employed. For more than two groups, one-way or two-way analysis of variance tests were applied, followed by posttests of either Tukey’s multiple comparisons or Fisher’s protected least significant difference tests. Statistical analyses were performed using from SAS Institute. The level of significance in all analyses was fixed at p < 0.05, as marked: *p < 0.05, **p < 0.01, ***p < 0.001, ****p < 0.0001

## Results

### ATF4 expression alters mitochondrial morphology in *Drosophila* flight muscle

To define mitochondrial organization across spatial scales, we combined Serial Block-Face Scanning Electron Microscopy (SBF-SEM) with confocal fluorescence imaging of intact tissue (Figure 1). High-resolution volumetric information on mitochondrial morphology was provided by SBF-SEM reconstructions (cyan), and assessment of mitochondrial alignment relative to the contractile apparatus was made possible by confocal imaging of mito-GFP (green) and phalloidin-FITC-labeled myofibrils (red). Figure 1A-C shows the results of three-dimensional SBF-SEM reconstructions revealing mitochondrial morphologies. Mitochondria in the control tissue were arranged in parallel sheets, parallel to the contraction axis, forming a dense, continuous network (Figure 1A). Rotated and orthogonal images (Figure 1A′, A″) showed a densely linked mitochondrial reticulum covering numerous sarcomeres, with limited vacant space and considerable mitochondrial connectivity. In contrast, mitochondria in the KD model exhibited pronounced structural disruption. The mitochondrial networks were found to be thin and perforated (Figure 1B), resulting in a porous, lattice-like structure. A loss of planar order and increased fragmentation were seen across the volume in the orthogonal views (Figure 1B′, B″). Figure 1C clearly shows a remodeling phenotype, with mitochondria looking stacked and expanded, creating thickened lamellar structures. In contrast to the controls, the rotated images (Figure 1C′, C″) exhibited reduced network connectivity but retained partial alignment with the contraction axis.

**Figure 1.**
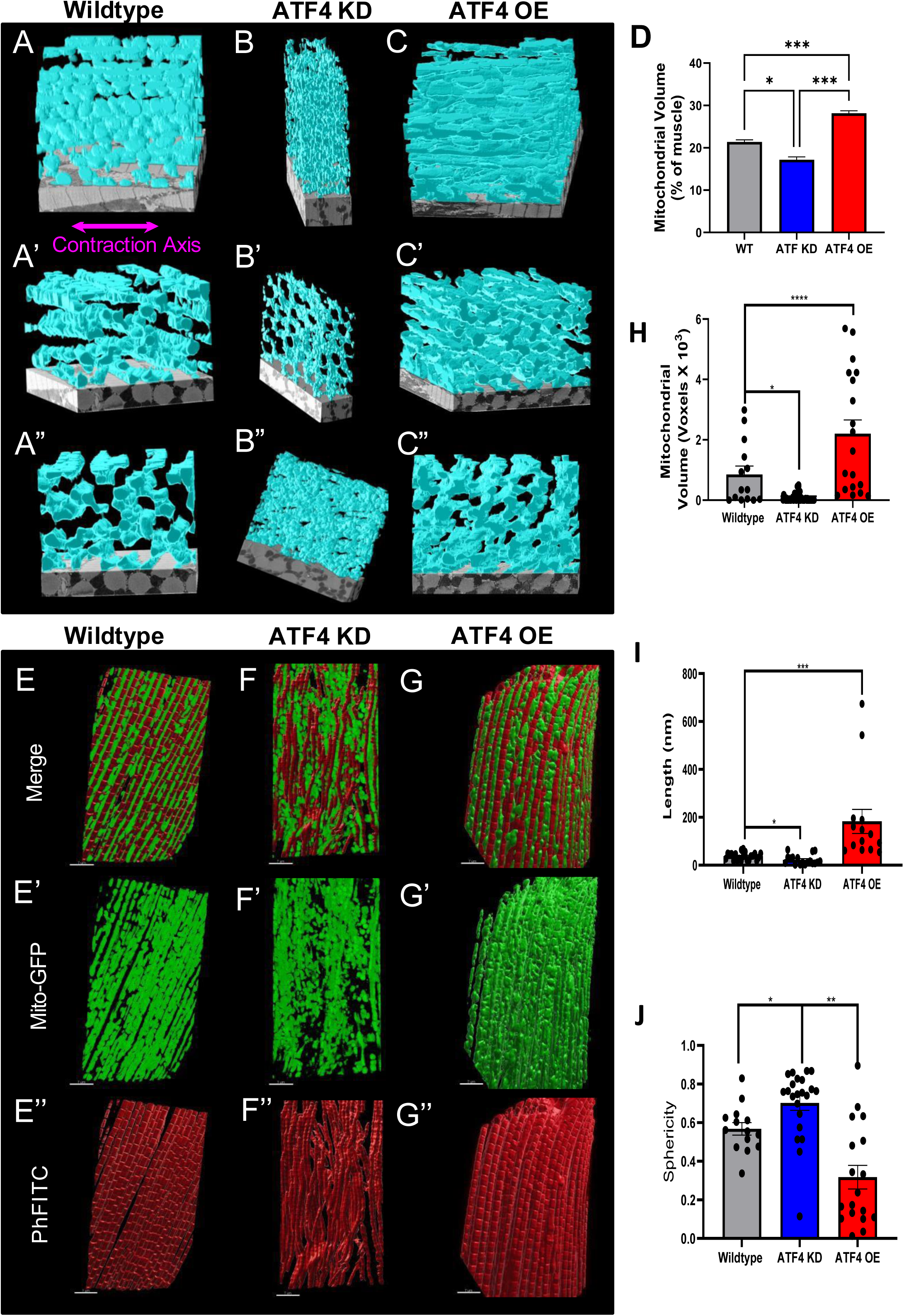
ATF4 expression alters mitochondrial morphology in *Drosophila* flight muscle. 3D reconstruction of Serial Block Face-Scanning Electron Microscopy (SBF-SEM) images for Wildtype (WT) (A-A”), ATF4 KD (B-B’”), and ATF4 OE (C-C”). Quantifications of mitochondrial volume between groups in A-C (D). 3D reconstruction using Imaris image analysis software of extracted flight muscle from WT (E), ATF4 KD (F), and ATF4 OE (G) models with GFP labeled mitochondria (E’, F’, and G’, respectively) and actin labeled in red (E”, F”, and G”, respectively). Quantitative comparisons between groups are provided for mitochondrial volume (H), mitochondrial length (I), and mitochondrial sphericity (J). Data was analyzed using an unpaired t-test. When more than two groups were compared, a one-way analysis of variance (ANOVA) was used, and statistical significance was determined by Tukey’s multiple comparisons test. A significant difference is defined as *p < 0.05, **p < 0.01, ***p < 0.001, ****p < 0.0001.

The SBF-SEM-identified ultrastructural characteristics were confirmed by confocal imaging (Figure 1E-G, Video1-12). To confirm the presence of planar mitochondrial sheets, mito-GFP-labeled mitochondria in control tissue (Figure 1E) organized into long, highly ordered arrays that closely tracked the direction of actin filaments. The mitochondrial architecture observed in the ATF4 KD group was characterized by a fragmented, punctate mito-GFP signal distributed unevenly across myofibrils in the disrupted samples (Figure 1F). Figure 1G shows that samples from the ATF4 OE groups exhibited expanded mitochondrial structures, with a thick, tightly aligned mito-GFP signal aligned with the contractile axis. The quantitative analysis of mitochondrial volume (Figure 1H) and length (Figure 1I) shows that they are increased in ATF4 OE and decreased in ATF4 KD, whereas sphericity (Figure 1J) is increased in ATF4 KD and decreased in ATF4 OE.

These findings indicate that genetic modulation of ATF4 expression induces distinct remodeling states characterized by fragmentation or hypertrophic stacking, both of which are detectable by confocal imaging but most clearly defined by volumetric electron microscopy. Collectively, these data validate a multiscale imaging approach for linking mitochondrial ultrastructure to mesoscale tissue organization.

### ATF4 expression alters mitochondria, MERCS, and gene expression in *Drosophila* flight muscle

Transmission electron microscopy (TEM), S/TEM, SBF-SEM, and confocal fluorescence imaging were utilized to determine the effects of ATF4 signaling on mitochondrial structure and connectivity in 2-dimensions versus the 3-dimensions examined in Figure 1 (Figure 2). Using this multiscale method, we were able to quantitatively evaluate the morphology of mitochondria, the organization at higher levels, and the connectivity mediated by nanotubes in different events.

**Figure 2.**
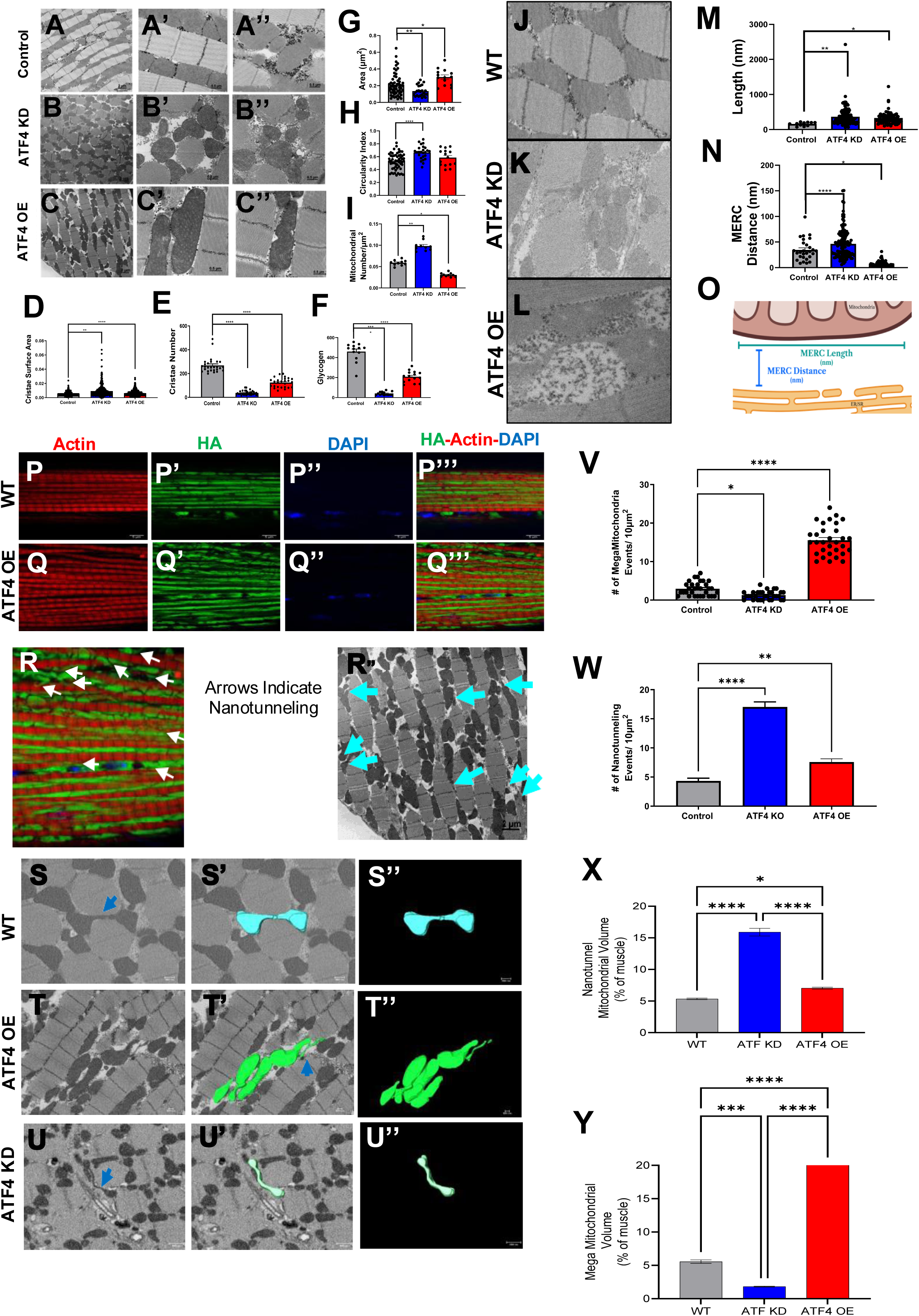
ATF4 expression alters mitochondria, MERCS, and gene expression in *Drosophila* flight muscle. TEM imaging shows alterations in WT. (A-A”), ATF4 KO (B-B”), and ATF4 OE (C-C”) mitochondria at different magnifications. Comparative quantifications between groups for cristae surface area (D), cristae number (E), muscle glycogen content as measured by glycogen-area following TEM imaging (F), mitochondrial area (G), mitochondrial circularity index (H), and mitochondrial number (I). Higher magnification TEM image of *Drosophila* flight muscle for WT (J), ATF4 KD (K), and ATF4 OE (L). Comparative quantification between groups for MERC length (M) and MERC distance (N). O) Representative schematic image showing the orientation of measurement for MERC distance and length. Confocal imaging of flight muscle from WT and ATF4 OE with actin labeled red (P and Q, respectively), HA labeled mitochondria (green) (P’ and Q’, respectively), DAPI stained nuclei (P” and Q”, respectively), and overlaid (P’’’ and Q’’’, respectively). Confocal (R) and TEM (R”) imaging of mitochondrial nanotunneling observed in ATF4 OE flight muscle as indicated by arrows. 3D reconstructions with orthoslice of nanotunnels in *Drosophila* flight muscle from SBF-SEM imaging for WT (S-S”), ATF4 OE (T-T”), and ATF4 KD (U-U”). TEM quantifications of Megamitochondria (V) and nanotunnel frequency (W) in control group, ATF4 OE, and ATF4 KD tissue. SBF-SEM datasets showing nanotunnels, mitochondrial volume and mega mitochondrial volume in control group, ATF4 OE, and ATF4 KD tissue. (a), mitochondrial dynamics (b), Ca^2+^ signaling (c), and Ca^2+^ transport (d). Data was analyzed using an unpaired t test. When more than two groups were compared, a one-way analysis of variance (ANOVA) was used, and statistical significance was determined by Tukey’s multiple comparisons test. A significant difference is defined as *p < 0.05, **p < 0.01, ***p < 0.001, ****p < 0.0001.

### ATF4 loss and overexpression induce distinct mitochondrial ultrastructural phenotypes

When ATF4 levels were altered, conventional TEM imaging showed significant changes in mitochondrial morphology with varying ATF4 expression levels in *Drosophila* indirect flight muscle (Figure 2A-C”). Mitochondria from our control group (Figure 2A-A″) were long and uniform in size, aligned with myofibrils, and displayed an undamaged cristae configuration. As seen in Figure 2B-B″, mitochondria from our ATF4 knockdown (ATF4 KD) group appear smaller and have a significantly higher circularity index (Figure 2 G, H), with increased cristae surface area and a decreased number of cristae, suggesting a possible disturbance in mitochondrial bioenergetics. Remarkably, mitochondria appear significantly larger and more elongated with ATF4 OE (Figure 2C-C″). This mitochondrial enlarged phenotype, or formation of Megeamitochondria, has been linked to increased metabolic function and stress-responsive mitochondrial remodeling; these mitochondria, which crossed numerous sarcomeres, displayed the characteristic enlarged volume and cross-sectional area previously seen by others with the use of chemically induced cellular stress models. The uneven arrangement of cristae inside these larger mitochondria, rather than a consistent enlargement, suggested structural adaptation taking place over time. Quantifications of these structural changes (Figure 2D-I) indicates that the size, shape, and density of the mitochondria evolved significantly in response to overexpression of ATF4. In contrast to ATF4 OE mitochondria, there was an increase in area and a decrease in aspect ratio in ATF4 KD mitochondria, suggesting that ATF4 plays a role in the maintenance of mitochondrial structural integrity, even without the presence of stress.

### SBF-SEM reveals altered mitochondria-ER contact sites (MERCS) with modulation of ATF4 expression

Next, we sought to determine whether the formation of MERCS was altered by manipulating ATF4 expression (Figure 2J-L). We found that although MERC length increases with both ATF4 KD (Figure 2 K) and OE (Figure 2 L) relative to control (Figure 2 J, M), MERC distance, however, significantly increased with ATF4 KD and significantly decreased with OE (Figure 2 J-L, N). These data indicate that although MERC length increases with loss of ATF4 and ATF4 OE, the distance between juxtaposed organelles increases with KD and decreases with OE, reflecting decreased and increased MERC tethering, respectively (Figure 2 J-O).

### Confocal imaging and SBF-SEM reveal an increased prevalence of Mitochondrial Nanotunnels with alterations in ATF4 expression

Throughout our investigation into how the expression and loss of ATF4 alters mitochondrial structure, we observed considerable alterations in the formation of thin, double-membrane bound protrusions that connect the matrix of the involved mitochondrion, called nanotunnels (Figure 2 P-U”, Video13-15). Specifically, the loss of ATF4 significantly alters mitochondrial morphology and produces distinct and highly organized, stereotypically textbook-like nanotunnel structures (Figure 2 U-U”). The thin mitochondrial protrusions observed with ATF4 KD resemble those described in metabolically restricted and stressed and may represent an adaptive effort to maintain organelle-organelle communication during impaired proteostasis and aberrant metabolic regulation. In contrast, the overexpression of ATF4 leads to the mitochondrial expansion and the formation of megamitochondria that remain contiguous through distinguishable nanotunnel-like bridges (Figure 2 P-Q’’’; a closer look with nanotunnels indicated by arrowheads shown in (R) and SBF-SEM images of nanotunnels shown in (R’)).Quantifications of these changes indicate that nanotunneling increases significantly with both ATF4 KD and OE compared with control flight muscle (Figure 2 P-Y). Although both the loss of and the overexpression of ATF4 relative to control lead to an increase in the prevalence of nanotunnels, these structures provide a means for the formation of extended mitochondrial networks through physical, elongated tubular connections joining adjacent mitochondria, suggesting enhanced content exchange and intra-mitochondrial communication.

Importantly, although these two nanotunnel phenotypes appear anatomically distinct, they may share essential functional similarities; both conditions likely reflect altered mitochondrial remodeling and stress states, potentially involving alterations in mitochondrial cristae structure, lipid metabolism, the distribution of mitochondrial DNA and the regulation of intra-organelle contacts. Thus, ATF4 levels may attune mitochondrial network plasticity, shifting between the compensatory formation of nanotunnels with loss-of-function and the hyperconnectivity of megamitochondria observed with gain-of-function.

### SBF-SEM reveals ATF4-dependent shifts in mitochondrial size and shape distributions

Mitotyping of individual mitochondria from wild-type, ATF4 KD, and ATF4 *Drosophila* OE indirect flight muscle (IFM) showed different physical characteristics dependent on ATF4 expression levels (Figure 3A). Highly oxidative flight muscle is characterized by a limited distribution of elongated, evenly sized mitochondria, which was observed in WT IFM mitochondria. On the other hand, mitochondrial morphologies in ATF4 KD IFM shifted toward smaller, rounder shapes (Figure 3A, F), suggesting disturbed mitochondrial homeostasis. Notably, ATF4 OE IFM showed a wide range of mitochondrial morphologies, including mitochondria that were significantly longer and wider and had a higher volume, which is compatible with mitochondrial expansion and the development of structures similar to Megamitochondria (Figure 3 A-F). These mitotyping results indicate that ATF4 expression significantly affects mitochondrial size, shape, and variability in *Drosophila* IFM.

**Figure 3.**
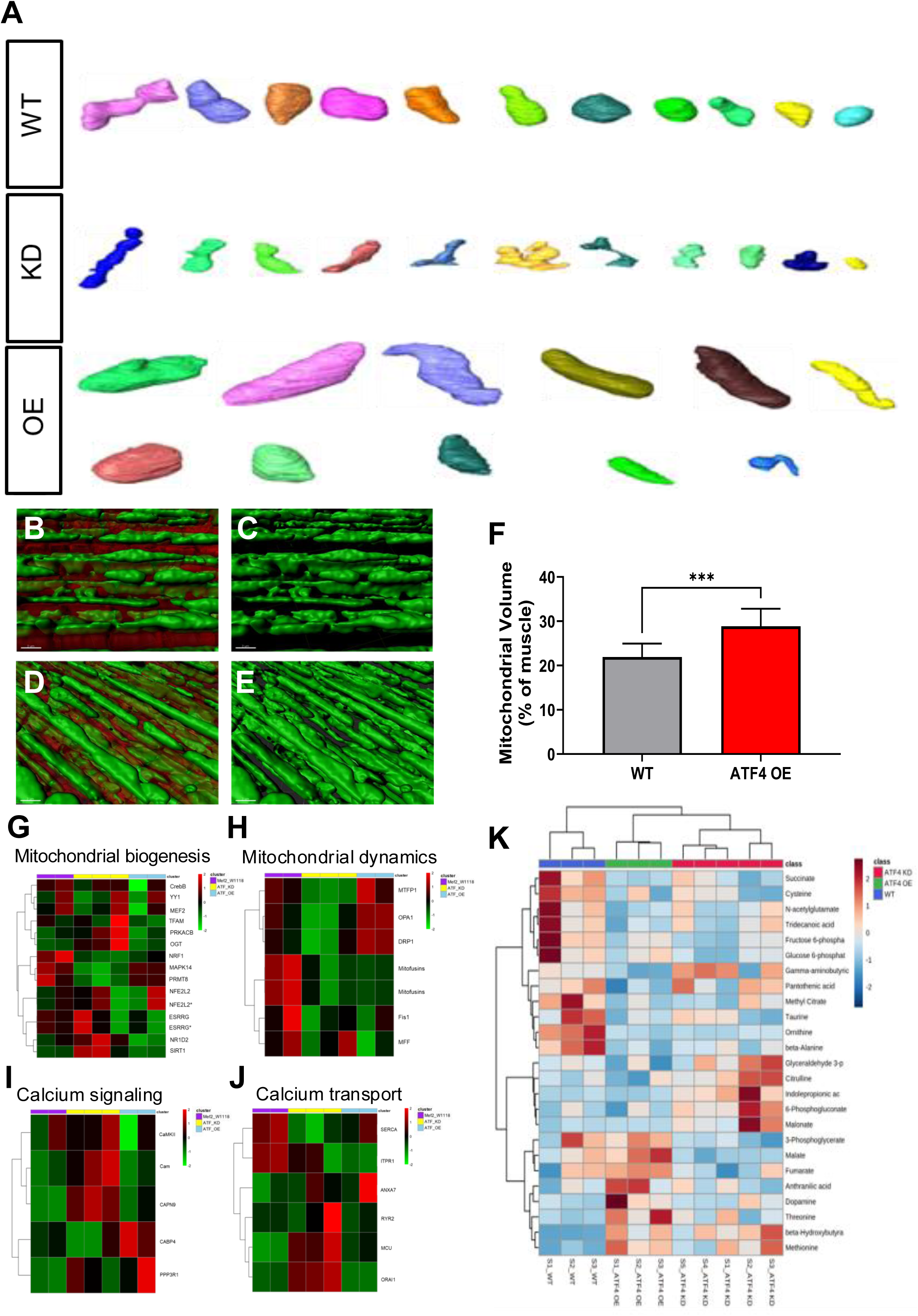
ATF4 expression modulates mitochondrial morphology in murine myotubes. Primary mouse myotubes were transduced with adenovirus expressing GFP (A), GFP-cre (B), or ATF4 OE (C) adenovirus. TEM imaging of control (D), ATF4 KO (E), ATF4 OE 24-hours post-infections (F), and ATF4 OE 48-hours post-infection (G). Comparative quantifications of mitochondrial area (H), cristae score (I), and MERC distance (J) between groups. Metabolomic profiling core metabolic pathways in IFM (K). Data was analyzed using an unpaired t test. When more than two groups were compared, a one-way analysis of variance (ANOVA) was used, and statistical significance was determined by Tukey’s multiple comparisons test. A significant difference is defined as *p < 0.05, **p < 0.01, ***p < 0.001, ****p < 0.0001.

### ATF4-dependent mitochondrial remodeling is coupled with transcriptional and metabolic reprogramming

After observing consistent mitochondrial morphology changes with manipulation of ATF4 expression, we sought to understand better the implications of these changes at both transcriptional (Figure 3 G-J) and metabolic (Figure 3 K) levels. We utilized RNA-sequencing to look at possible changes at the transcriptome level with alterations in ATF4 expression. According to heatmap analysis, genes related to mitochondrial dynamics, oxidative metabolism, stress-responsive signaling, and quality control pathways showed coordinated expression changes in ATF4 OE. Additional evidence from metabolomic profiling showed that ATF4-dependent remodeling of core metabolic pathways also occurred (Figure 3K). There were changes in metabolites linked to glycolysis, the tricarboxylic acid cycle, amino acid metabolism, and redox balance in ATF4 OE samples, which separated them from WT and ATF4 KO samples. These alterations are in line with the hypothesis that ATF4 expression in *Drosophila* IFM leads to increased mitochondrial mass and a response to different energy demands.

### ATF4 links mitochondrial architecture to calcium homeostasis in myotubes

To examine the functional implications of our observed mitochondrial morphology and nanotunnel changes in a mammalian system, we examined primary mouse WT, ATF4 KO, and ATF4 OE myotubes from ATF4^fl/fl^ mice (Figure 4). Using fluorescent (Figure 4 A-C) and TEM (Figure 4 D-G) imaging, we consistently observed successful infection with adenovirus expressing GFP (control; A), cre-recombinase with a GFP reporter (ATF4 KO; B), and ATF4 adenovirus with a GFP reporter (C). Looking at the mitochondrial level via TEM images, we found that a loss of ATF4 resulted in significant decreases in mitochondrial area and cristae score (or the quality of cristae present), with concomitant increases in MERC distance, suggesting a loss or decrease in MERC tethering relative to control (Figure 4D, E, H-J). Alternatively, the overexpression of ATF4 resulted in significant increases in mitochondrial area and cristae score relative to control and concomitant decreases in MERC distance, indicating an increase in MERC tethering (Figure 4 F-J). To better understand how these MERC changes might alter calcium homeostasis, we used the radiometric dye, Fura-2 AM, to measure cytosolic calcium dynamics (Figure 4K-M). In all situations, stimulation caused cytosolic calcium levels to spike quickly, but the response’s intensity and rate of elevation were heavily dependent on ATF4 (Figure 4K). Peak calcium amplitude quantification showed that ATF4-deficient and ATF4 OE myotubes had significantly lower peak cytosolic calcium levels than the control group (Figure 4K, L). Area under the curve (AUC) analysis showed that cumulative calcium signal was much higher in ATF4 deletion compared to control, but ATF4 overexpression was not significantly different from control (Figure 4M), suggesting that ATF4 absence altered buffering and calcium clearance dynamics. Taken together, these data indicate that ATF4 expression alters mitochondrial size, MERC distance, and cytosolic calcium handling in mouse primary myotubes.

**Figure 4.**
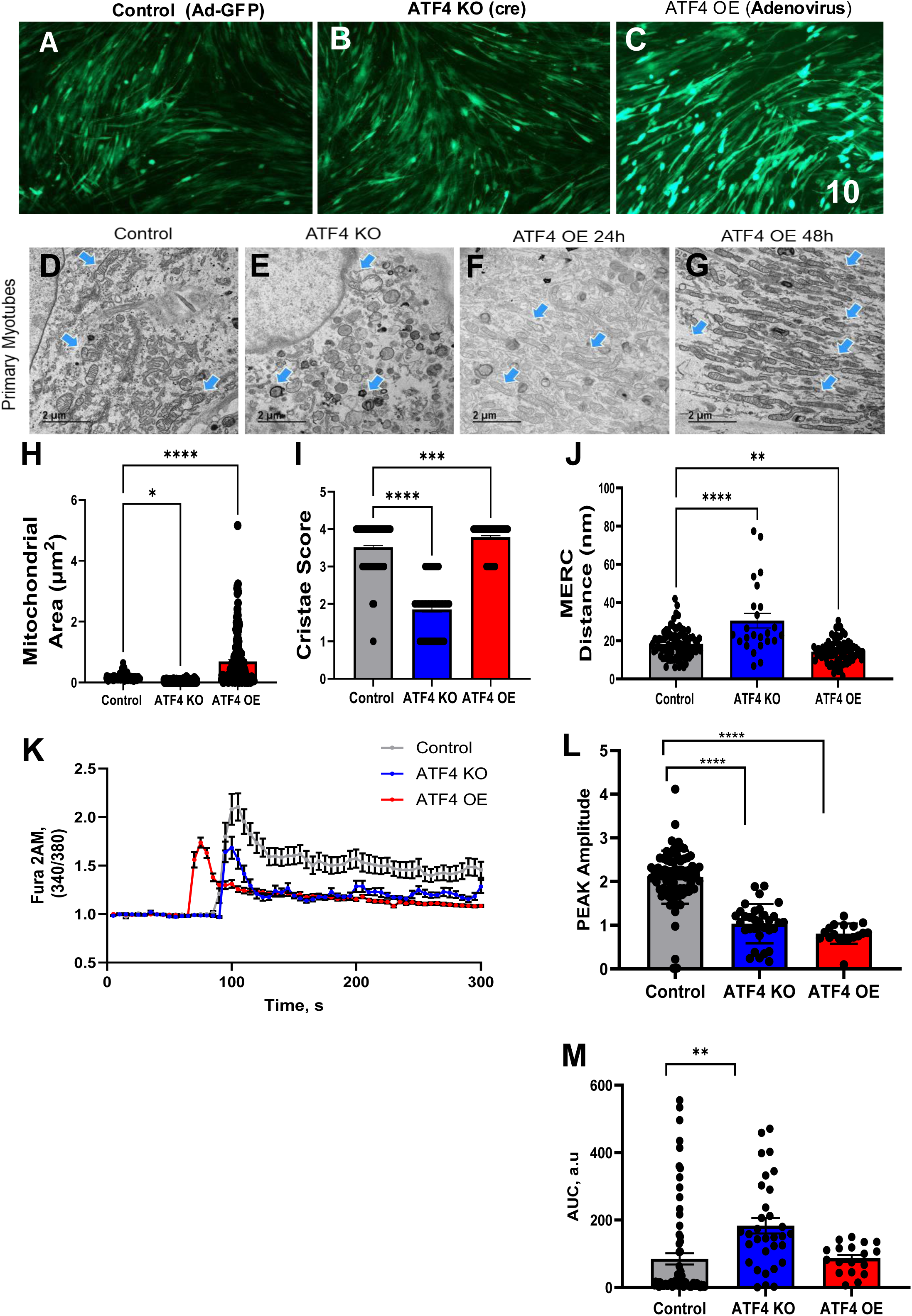
ATF4 regulates mitochondrial ultrastructure and calcium handling in myotubes. (A–C) Representative fluorescence images of myotubes infected with either adenovirus expressing GFP or ATF4. The presence of fluorescence indicates successful infection. Scale bar, 10X. (D–G) Transmission electron microscopy (TEM) images of myotubes reveal ultrastructural mitochondrial remodeling associated with ATF4 expression. Control myotubes exhibit elongated mitochondria with well-organized cristae aligned along the contractile axis. In contrast, ATF4 manipulation results in altered mitochondrial size, shape, and cristae organization, with increased mitochondrial heterogeneity and disruption of longitudinal alignment. Blue arrows indicate representative mitochondria. Scale bar, 2 µm. (H–J) Quantification of mitochondrial ultrastructural parameters in myotubes, including mitochondrial area, aspect ratio, and cristae density. ATF4 overexpression (OE) and ATF4 knockout (KO) differentially regulate mitochondrial morphology, consistent with distinct effects on mitochondrial fusion–fission balance and internal membrane organization. Data are presented as mean ± SEM. (K) Representative cytosolic calcium traces measured by Fura-2 AM (340/380 nm) in myotubes following stimulation. Control myotubes display robust and sustained calcium transients, whereas ATF4 KO and ATF4 OE myotubes exhibit blunted calcium responses, indicating impaired calcium handling. (L) Quantification of peak calcium amplitude reveals a significant reduction in calcium signaling in ATF4 KO and ATF4 OE myotubes compared with controls. ****P < 0.0001. (M) Area under the curve (AUC) analysis of calcium transients demonstrates reduced overall calcium signaling in ATF4 KO myotubes compared with controls, whereas ATF4 OE shows no significant difference relative to control. **P < 0.01; ns, not significant.

### SBF-SEM reveals ATF4-dependent remodeling of MERC volume and length in 3D

To determine if the MERC changes we observed in our mammalian system for control, ATF4 KO, and ATF4 OE in 2-dimensions were consistent in 3-dimensions, we utilized SBF-SEM reconstructions (Figure 5A-C’). Similar to our observed results in 2-dimensions, ATF4 KO showed significantly decreased MERC volume and MERC length (Figure 5 B-B’, D, E) relative to control (Figure 5 A-A’, D, E) In contrast to our ATF4 KO model, but consistent with our 2D findings, ATF4 OE resulted in significantly increased MERC volume and MERC length when compared to control (Figure 5 A-A’, C-C’, D, E). These findings are consistent with previous reports that MERC tethering is ATF4-dependent in OPA1-deficient myotubes (Hinton, Katti et al. 2024).

**Figure 5.**
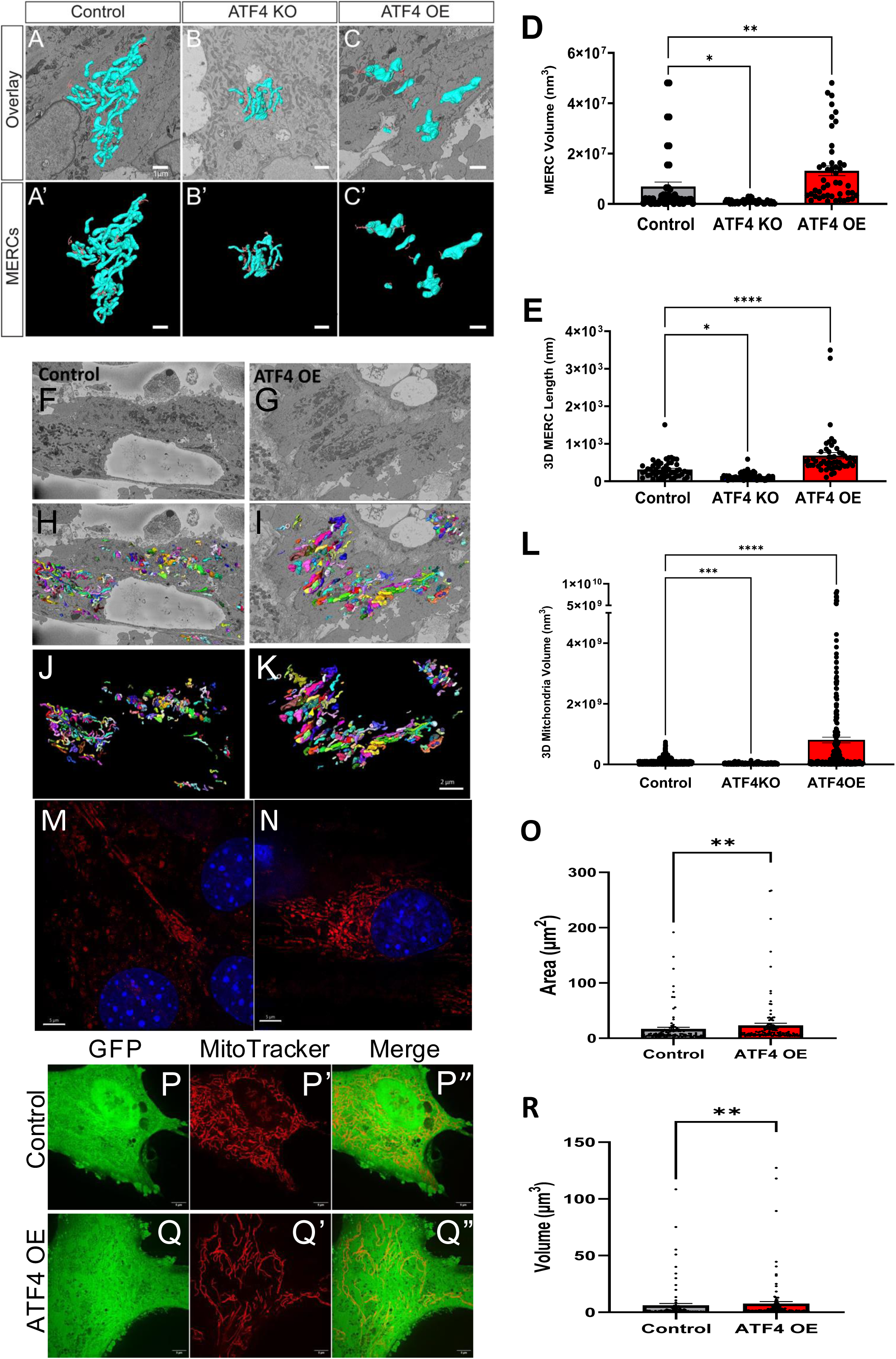
ATF4 regulates mitochondrial–ER contact architecture and mitochondrial network organization in myotubes and myoblasts. (A–C) Representative serial block-face scanning electron microscopy (SBF-SEM) images of myotubes showing mitochondrial–ER contact sites (MERCS; cyan) overlaid on ultrastructural images under Control, ATF4 knockout (KO), and ATF4 overexpression (OE) conditions. (A – C) Three-dimensional reconstructions of MERCS segmented from SBF-SEM datasets corresponding to panels A–C, illustrating ATF4-dependent changes in MERC size, continuity, and spatial organization. (D) Quantification of total MERC volume (nm³) per myotube demonstrates a significant increase in MERC volume with ATF4 OE and a marked reduction with ATF4 KO compared with Control. (E) Quantification of total 3D MERC length (nm) per myotube confirms ATF4-dependent enhancement of ER–mitochondria contact length, with ATF4 OE showing increased MERC length and ATF4 KO showing reduced contacts. (F–G) Representative transmission electron microscopy (TEM) images of myotubes highlighting longitudinal mitochondrial alignment and ER–mitochondria interfaces in Control and ATF4 OE conditions. (H–I) Pseudocolored segmentation of mitochondria from TEM images in panels F–G, illustrating increased mitochondrial elongation and network complexity in ATF4 OE myotubes. (J–K) Three-dimensional reconstructions of mitochondrial networks in myotubes reveal increased mitochondrial volume and connectivity in ATF4 OE relative to Control. (L) Quantification of total mitochondrial volume (nm³) in myotubes shows a significant increase in mitochondrial volume with ATF4 OE compared with ATF4-deficient myotubes. (M–N) Representative stimulated emission depletion (STED) microscopy images of C2C12 myotubes stained for mitochondria (red) and nuclei (blue), demonstrating altered mitochondrial network density and perinuclear organization under ATF4 manipulation. (P) Spinning disk confocal images of control primary myoblasts expressing GFP and stained with MitoTracker. (P) GFP (green) marks the cell body; (P) MitoTracker (red) labels the mitochondrial network; (P) merged image illustrates mitochondrial distribution relative to cell morphology. (Q) Spinning disk confocal images of ATF4 OE myoblasts expressing GFP and stained with MitoTracker. (Q) GFP (green); (Q) MitoTracker (red); (Q) merged image reveals increased mitochondrial network density and elongation in ATF4 OE primary mouse myoblasts compared with control.

### ATF4 overexpression increases mitochondrial volume in primary myotubes

To assess if mitochondrial parameters changed in a consistent manner with MERC changes, we utilized SBF-SEM reconstructions for volumetric analysis (Figure 5 F-L) from ATF4^fl/fl^ primary mouse myotubes. Consistent with our previous findings, we found that ATF4 OE myotubes had a significantly larger mitochondrial volume when compared to (Figure 5 F-L). These findings are also consistent with the appearance of larger structures resembling Megamitochondria and indicate that the increase is due to mitochondrial expansion rather than simple redistribution. Taken together, these results show that ATF4 controls the number and architecture of mitochondria and the mitochondria-ER interface in a coordinated manner.

To look at these mitochondrial changes at a higher resolution we utilized STED super-resolution imaging in C2C12 myotubes (Figure M-N). Quantifications of these findings show that both mitochondrial area and volume significantly increase with ATF4 overexpression (Figure O, R). To look at these changes in a live-cell model, we utilized spinning-disk confocal SORA imaging of primary mouse myoblasts (Figure P-Q”). The mitochondrial networks in ATF4 OE myotubes appear more robust and elongated or hyperfused as indicated by MitoTracker (Figure P’, Q’). Cumulatively, these data show that ATF4-dependent mitochondrial remodeling is robust in C2C12 and primary mouse myoblasts and myotubes and that these findings are readily observable using live-cell imaging.

### ATF4 expression alters genes associated with mitochondria and MERCS in *Drosophila* flight muscle

To define the transcriptional programs downstream of ATF4 that may underlie the structural and functional mitochondrial phenotypes observed in vivo, we performed untargeted RNA sequencing on *Drosophila* muscle under ATF4 loss- and gain-of-function conditions (Figure 6).

**Figure 6.**
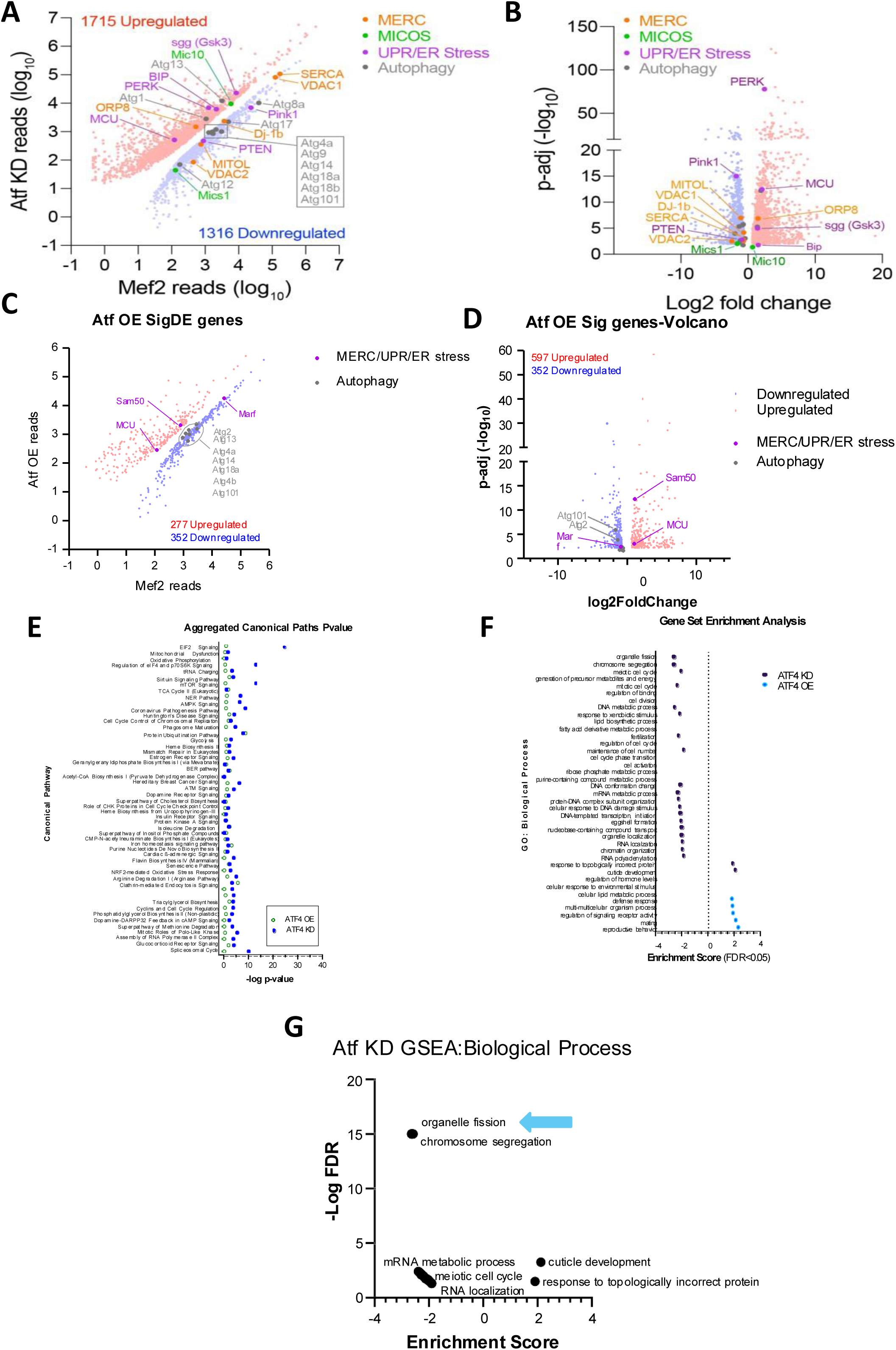
ATF4 expression alters genes associated with mitochondria and MERCs in *Drosophila* flight muscle. A) RNA sequencing data from ATF4 KD showing differentially expressed genes. B) Volcano plot of p-adjusted RNA sequencing data shown in (A). C) RNA sequencing data from ATF4 OE showing differentially expressed genes. D) Volcano plot of p-adjusted RNA sequencing data shown in (C). E) Changes in canonical pathways from RNA sequencing data from ATF4 KD and OE. F) Differentially expressed upstream regulators from RNA sequencing data from ATF4 KD and OE. G) Gene set enrichment analysis (GSEA) for biological processes in ATF4 KD and OE flight muscle. H)

#### ATF4 loss induces widespread transcriptional reprogramming in fly muscle

Figure 6 A-B shows the results of a comparative analysis between ATF4 KD and control muscles, which showed significant differences in transcription. Analysis of scatter plots comparing the ATF4 KO and control groups showed that gene expression shifted widely after ATF4 deletion, with 1,715 genes upregulated and 1,316 downregulated (Figure 6A). In a coordinated manner, genes related to mitochondrial structure and organelle communication, such as the MERC protein machinery and MICOS complex components, were downregulated, as determined by differential expression analysis. Volcano plot analysis provided additional evidence that transcripts related to mitochondrial dynamics, calcium handling, and organelle stress responses are significantly reduced in ATF4 KO muscle (Figure 6B). It has been observed that genes related to mitochondrial quality control, MICOS organization, and MERCS were inversely affected, indicating that ATF4 is necessary for the maintenance of transcriptional programs that promote mitochondrial architecture and organelle-organelle communication.

#### ATF4 overexpression activates stress-responsive and organelle remodeling pathways

Alternatively, a different transcriptional signature was observed in fly muscle with ATF4 OE (Figure 6C-D). Transcripts linked to autophagy-related pathways, unfolded protein response (UPR/ER stress), and mitochondria-ER interactions were significantly overrepresented in the upregulated genes identified by scatter plot analysis (Figure 6C). The volcano plots demonstrated strong activation of canonical ATF4 target pathways, which encompass genes involved in ER stress signaling, adaptive metabolic remodeling, and mitochondrial calcium handling (Figure 6D). Crucially, ATF4 OE muscle showed a specific upregulation of many MERC- and MICOS-associated genes, which allowed us to understand the transcriptional basis of the mitochondrial remodeling and increased MERC architecture we observed with SBF-SEM, STED, and confocal imaging.

#### Aggregated pathway and gene set enrichment analyses identify ATF4 as a central regulator of mitochondrial–ER communication networks

Figure 6E-G shows differential analysis reports for the ATF4 KO and ATF4 OE groups. These findings indicate that ATF4 regulates genes associated with mitochondrial structure, organelle-organelle contact formation, stress adaptation, and quality control. ATF4 acts as a transcriptional regulator, that modulates mitochondrial-ER communication, and stress-responsive remodeling is supported by the reciprocal regulation of genes related to MERCS, MICOS organization, calcium transport, and autophagy in response to ATF4 status. In line with the reported phenotypes of altered MERC size, mitochondrial volume expansion, and calcium-handling abnormalities, pathway-level analysis revealed coordinated modulation of the mitochondrial calcium uptake machinery, endoplasmic reticulum stress sensors, and genes associated with autophagy. These RNA-seq results show that ATF4 is a master transcriptional regulator in fly muscle, linking the biology of organelle contact sites, mitochondrial architecture, and cellular stress adaptation.

### ATF4 expression reciprocally alters mitochondrial bioenergetics and gene expression levels in primary mouse myotubes

To better understand the energetic functional implications at the mitochondrial level of ATF4 expression modulation, we utilized untargeted metabolomics and Seahorse extracellular flux analysis to measure oxygen consumption rates (OCR) in primary mouse myotubes. We found that the control, ATF4 KO, and ATF4 OE groups exhibit different metabolic states dependent on the expression of ATF4 (Figure 7A). The metabolic pathways related to amino acids, carbohydrates, and mitochondria were shown to undergo extensive alterations, as shown by differential metabolite analysis (Figure 7B).

**Figure 7.**
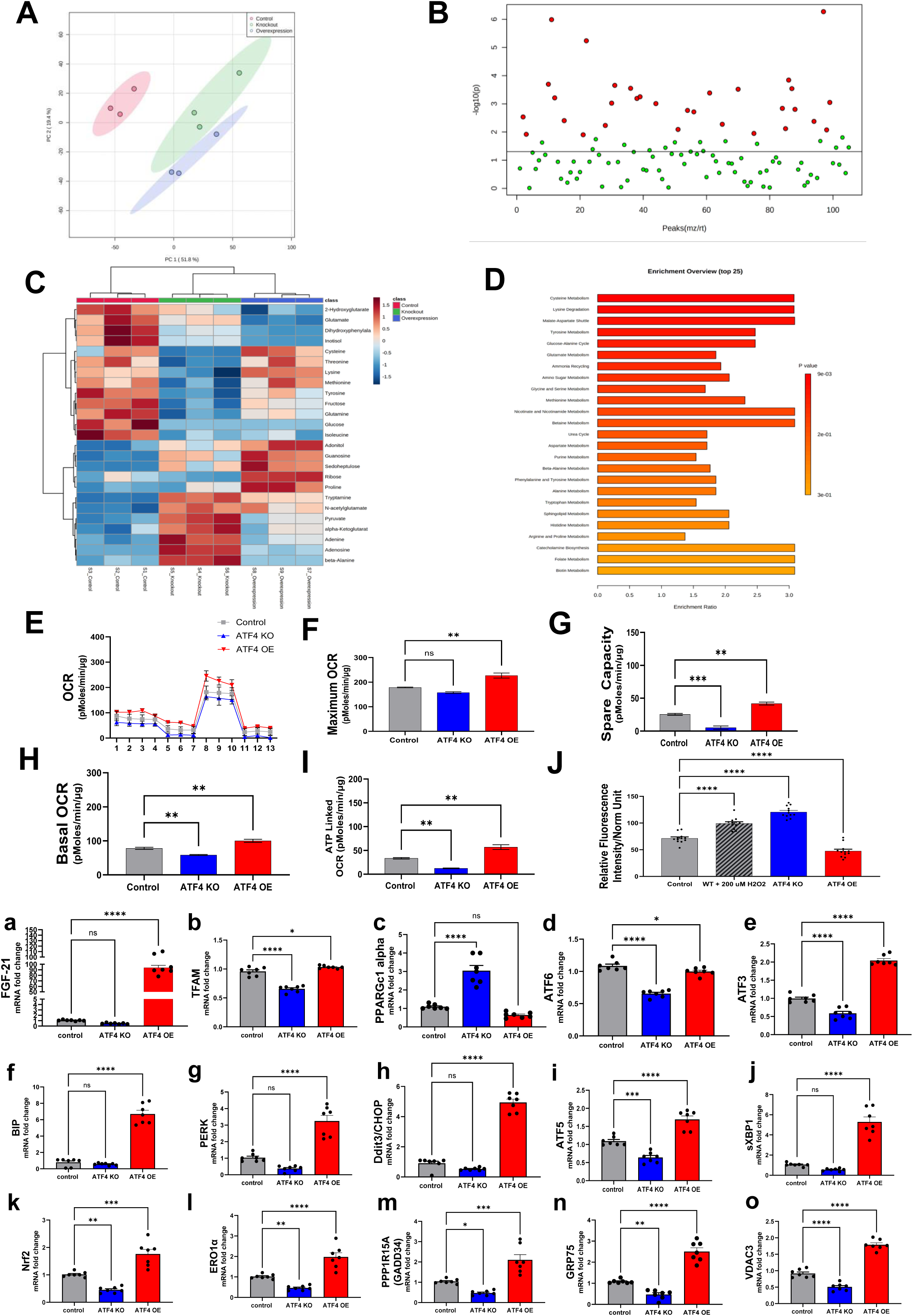
ATF4 overexpression alters mitochondrial function and gene expression levels. A) Principal component analysis (PCA) score plot based on primary mouse myotubes of Control, ATF4 KO, and ATF4 OE models (*n* = 3). B) Bubble plot of fold change abundance of metabolic species in Control (Red) and ATF4 KO (Green) primary mouse myotubes. C) Differential heatmap from metabolomic data for control, ATF4 KO, and ATF4 OE primary mouse myotubes. H) Enrichment analysis from the obtained metabolomic data obtained from each group from primary mouse myotubes. I) Oxygen consumption rate (OCR) measurements from each group. Comparative quantifications for maximum OCR (J), spare capacity (K), basal OCR (L), ATP-linked OCR (M), and mitochondrial H2O2 content (N) between groups measured by mitochondrial peroxy yellow 1 (MitoPY1). Relative gene expression levels between groups for FGF21 (a), TFAM (b), PPARGc1 (c), PGC1 alpha (d), ATF3 (e), BIP (f), PERK (g), CHOP (h), ATF5 (i), sXBP1 (j), NRF2 (k), ERO1α (l) GADD34 (m), GRP75 (n) VDAC3 (o) from primary mouse myotubes. Data was analyzed using an unpaired t-test. When more than two groups were compared, a one-way analysis of variance (ANOVA) was used, and statistical significance was determined by Tukey’s multiple comparisons test. A significant difference is defined as *p < 0.05, **p < 0.01, ***p < 0.001, ****p < 0.0001.

Metabolites associated with redox, glycolytic intermediates, and the tricarboxylic acid (TCA) cycle were among those that underwent alterations, as shown by heatmap visualization of significantly changed metabolites (Figure 7C). Additionally, metabolic pathways linked to amino acid biosynthesis, stress-responsive metabolic adaptability, and mitochondrial energy metabolism were shown to be significantly overrepresented in the pathway enrichment analysis (Figure 7D). Using a Seahorse extracellular flow analysis to assess if metabolic reprogramming resulted in functional mitochondrial alterations. In control myotubes, ATF4 overexpression greatly enhanced baseline and maximal oxygen consumption rates, whereas ATF4 deletion diminished respiratory capacity (Figure 7E-G). The results show that ATF4 increases mitochondrial oxidative capacity, which is consistent with the expanded organelle networks and larger mitochondrial volume observed in EM and super-resolution imaging. Further investigation into spare respiratory capacity revealed that ATF4 OE myotubes maintain greater bioenergetic flexibility than ATF4-deficient cells, which exhibit a diminished ability to meet increased energetic demand. Together, these data indicate that ATF4 drives both metabolic reprogramming and mitochondrial bioenergetic capacity in primary mouse myotubes.

#### ATF4 regulates cellular redox state and reactive oxygen species (ROS) production

We next sought to understand the functional implications of altered mitochondrial energetics with ATF4 expression by measuring intracellular ROS levels. We observed an increase in ROS buildup following ATF4 deletion, consistent with altered calcium homeostasis and reduced mitochondrial activity (Figure 7J). On the other hand, ATF4 overexpression kept ROS levels stable or slightly decreased them even though mitochondrial respiration increased (Figure 7 H-J), indicating that ATF4 limits oxidative stress by coordinating mitochondrial expansion with redox regulatory systems.

#### ATF4-dependent metabolic remodeling is supported by transcriptional changes

We utilized targeted qPCR analysis (Figure 7a–o) to examine whether metabolic and functional alterations were occurring with alterations in mitochondrial and metabolic gene transcription levels. Overexpression of ATF4 increased expression of genes regulating mitochondrial metabolism, stress adaptation, and redox homeostasis, whereas ATF4 deletion reduced their expression. These results from metabolomic and bioenergetic studies corroborate the idea that ATF4 regulates metabolic pathways by transcriptional control to facilitate mitochondrial growth, improve respiration, and maintain redox equilibrium. Collectively, our data indicate that loss of ATF4 leads to metabolic dysregulation, reduced respiration, and elevated oxidative stress, whereas ATF4 overexpression causes metabolic reprogramming, improves mitochondrial respiratory capacity, and preserves redox balance.

### Induced stress increases ATF4 expression and MERC tethering in human and mouse myotube

To gain deeper insight into ATF4 induction during cellular stress, we utilized oligomycin, an inhibitor of mitochondrial ATP synthase, to induce cellular stress in primary human and mouse myotubes. Western blot analysis showed that ATF4 protein expression levels increased with time in primary mouse (Figure 8A) and primary human (Figure 8 B) myotubes. Quantifications of these changes and the protein level changes in MERC tethering machinery (Figure 8 D-H) with induced cellular stress indicate mitochondrial-organelle communication responses initiate quickly and remain elevated with prolonged oligomycin-induced cellular stress.

**Figure 8.**
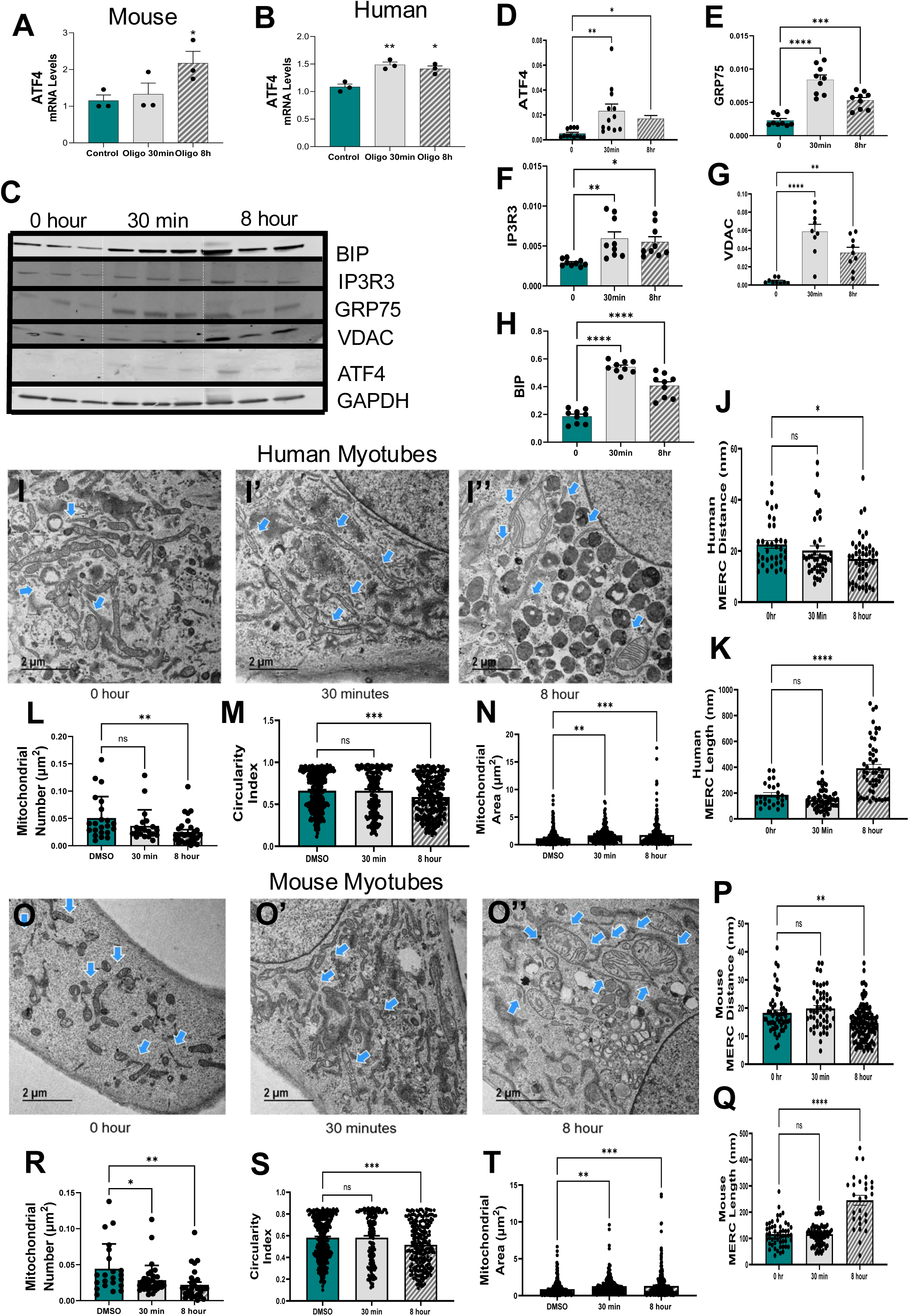
Induced stress increases ATF4 expression and MERC tethering. Relative gene expression of ATF4 following oligomycin treatment for 0-hour, 30-minute and 8-hour in primary mouse (A) and primary human (B) myotubes. C) Representative Western blot and quantifications for ATF4 (D) and MERC tethering proteins: GRP75 (E), IP3R3 (F), VDAC (G), and BIP (H) following oligomycin treatment in primary mouse myotubes. I) TEM imaging of human primary myotubes treated with Oligomycin for 0-minutes, 30-minutes (I’), and 8-hours (I”). Quantitative comparisons for MERC distance (J), MERC length (K), mitochondrial number (L), mitochondrial circularity index (M), and mitochondrial area (N) changes between groups shown in I-I” following Oligomycin treatment. TEM imaging of primary mouse myotubes treated with Oligomycin for 0-minutes (O), 30-minutes (O’), and 8-hours (O”). Quantitative comparison of MERC distance (P), MERC length (Q), mitochondrial number (R), mitochondrial circularity index (S), and mitochondrial area (T) changes in primary mouse myotubes following—Oligomycin—Oligomycin treatment. Data was analyzed using an unpaired t test. When more than two groups were compared, a one-way analysis of variance (ANOVA) was used, and statistical significance was determined by Tukey’s multiple comparisons test. A significant difference is defined as *p < 0.05, **p < 0.01, ***p < 0.001, ****p < 0.0001.

#### TEM analysis reveals increased Megamitochondria formation and MERC tethering with oligomycin induced cellular stress

To assess if the increases in ATF4 and MERC tethering machinery protein expression occur concomitantly with the formation of Megamitochondria, we utilized TEM imaging of primary mouse and human myotubes (Figure 8 I-T). We found that prolonged treatment with oligomycin significantly decreased mitochondrial number while simultaneously increasing mitochondrial area in human (Figure 8 I-I”, L-N) and mouse (Figure 8 O-O”, R-T) primary myotubes. We also found significant decreases in MERC distance after 8-hours of oligomycin treatment (Figure 8 I-I”, J, K, O-O”, P, Q) with significant increases in MERC length, indicating increased tethering and communication in both human (Figure 8 I-I”, J, K) and mouse (Figure 8 O-O”, P, Q) primary myotubes. These data collectively indicate that inducing cellular stress with oligomycin treatment produces results in human and mouse myotubes that are consistent with our ATF4 overexpression model.

### ATF4-driven mitochondrial remodeling is dependent on MFN2 in myotubes

Since mitochondrial morphology is driven and determined by mitochondrial dynamic, we want to determine if the ATF4-induced mitochondrial remodeling we observe is dependent on mitochondrial dynamics machinery. To investigate this, we first measured the abundance of proteins involved in mitochondrial dynamics in ATF4 OE cells. Western blot analysis revealed that with ATF4 overexpression, MFN2 is the only mitochonchondrial dynamics protein that is significantly increased (Figure 9 A-H).

**Figure 9.**
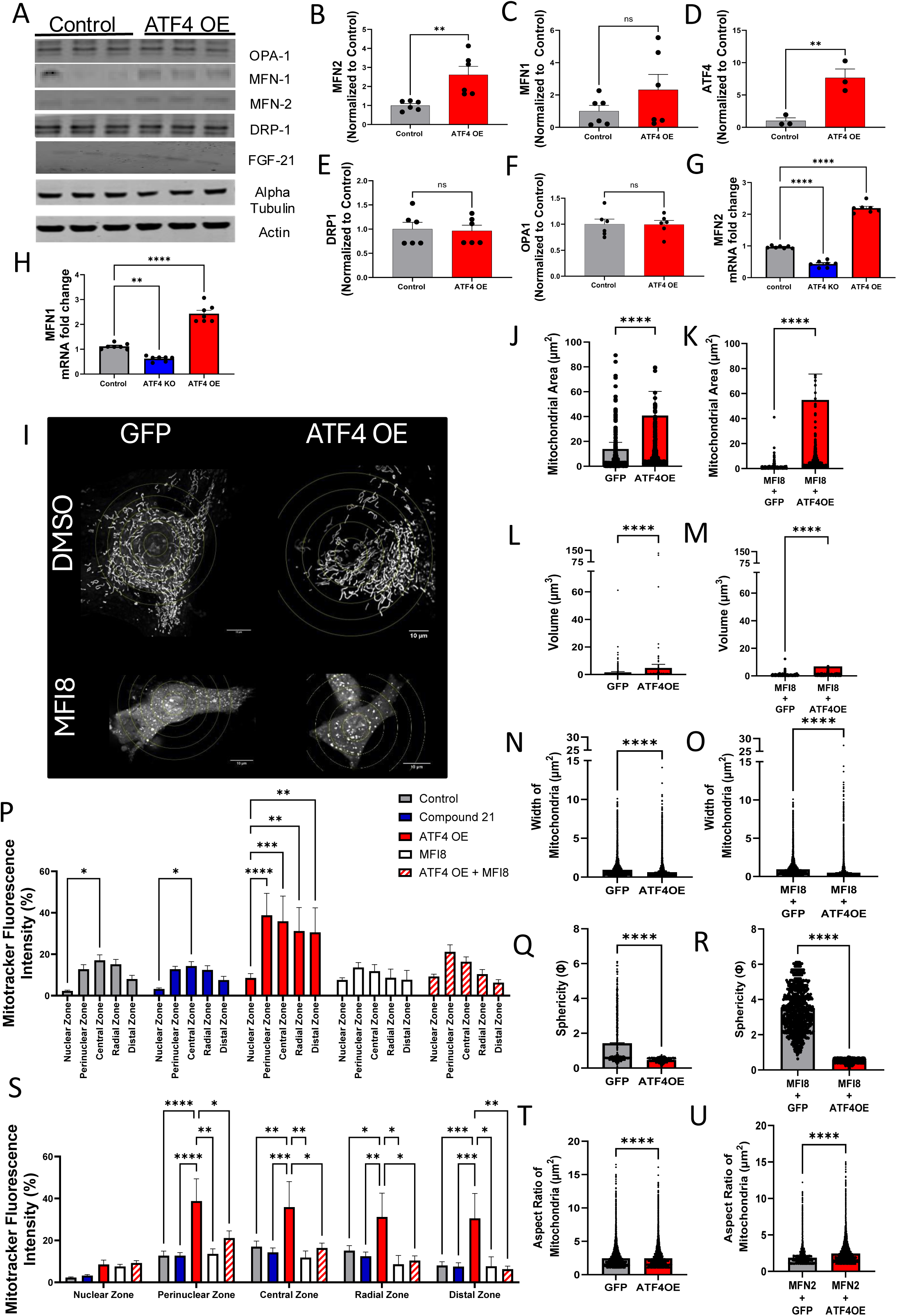
Expression levels for proteins involved in mitochondrial dynamics change with ATF4 expression. A) Representative Western blots from control and ATF4 OE primary myotubes showing mitochondrial dynamic proteins. Comparative quantifications of protein levels for MFN2 (B), MFN1 (C), ATF4 (D), DRP1 (E), and OPA1 (F). Comparison of relative gene expression levels for MFN2 (G) and MFN1 (H) in primary mouse myotubes between Control, ATF4 KO, and ATF4 OE. I) Representative confocal microscopy images revealing mitochondrial morphology for GFP Control or ATF4 OE cells treated with dimethyl sulfoxide (DMSO) or the MFN2 oligomerization inhibitor, MFI8. J, K) Mitochondrial area (J) and aspect ratio (K) intensity analyses under basal conditions for DMSO or MFI8 treatments, demonstrating that ATF4 OE increases mitochondrial size and elongation through MFN2. L–O) Histograms for mitochondrial area (L, M) and aspect ratio (N, O) comparing Control versus ATF4 OE cells. P) Analysis of mitochondrial network features under all experimental conditions. Q, R) Percentage of total network versus separates mitochondria for Control and ATF4 OE cells. S) Mitochondrial network measures for all treatment groups. T, U) Additional mitochondrial morphology studies under Control versus ATF4 OE conditions. Data represent mean ± SEM. *P < 0.05, **P < 0.01, ***P < 0.001, ****P < 0.0001.

#### ATF4-induced mitochondrial network expansion is sensitive to MFN2 inhibition

To determine whether MFN2 was required for the mitochondrial changes observed with ATF4 OE, we used a chemical inhibitor of mitochondrial oligomerization, FMI8. Using spinning disc confocal microscopy, we observed significant changes in MFI8-treated primary myoblasts, both with and without ATF4 overexpression (Figure 9I). The results show that ATF4 OE-induced requires functional MFN2-mediated fusion.

Consistent with our other data, quantifications of mitochondrial parameters in our live-cell imaging experiments reveal increases in mitochondrial area and volume and decreased mitochondrial sphericity with ATF4 OE (Figure 9 J, L). Notably, although the addition of MFI8 with ATF4 OE shows significant increases in both mitochondrial area and volume and significant decreases in mitochondrial sphericity in comparison to MFI8 treated GFP control, both groups showed significant mitochondrial fragmentation (Figure 9 K, M).

#### MFN2 inhibition uncouples ATF4 signaling from mitochondrial expansion

Collectively, these results show that ATF4 overexpression stimulates mitochondrial network growth via a process that depends on fusion and involves MFN2. The fact that ATF4-induced mitochondrial remodeling can be blocked by pharmacological suppression of MFN2 oligomerization even when ATF4 expression is sustained suggests that MFN2 is located downstream of ATF4 in the pathway that regulates mitochondrial architecture. Concentric circle analysis revealed that suppression of MFN2 oligomerization prevents the mitochondrial redistribution observed with ATF4 overexpression (Figure 9P, S). According to zone-based quantification, ATF4 whereas blocking MFN2 oligomerization prevents this redistribution. Taken together, these results show that ATF4 transcriptionally programs a coordinated program that leads to increased mitochondrial fusion and elongation. Together, these findings indicate that MFN2 is a crucial effector of ATF4-driven mitochondrial remodeling (Fig. 9 T–U).

### ATF4 binds to the promotors of NRF1 and Nrf2 during cellular stress to facilitate MFN2-dependent mitochondrial remodeling in C2C12 myoblasts

Since it has previously been reported that ATF4 does not bind the promoter of the MFN2 gene, we first confirmed these findings using ChIP-seq in tunicamycin-treated mouse embryonic fibroblasts (MEFs) (Supplemental Figure). Since our findings are consistent with previous reports that ATF4 does not bind to the *Mfn2* promoter directly under tunicamycin-induced cellular stress, we then considered other transcription factors known to bind the Mfn2 promoter directly. Interestingly, we found that ATF4 binds to the promoters of the genes encoding NRF1 and Nrf2 during cellular stress, both of which are known to control MFN2 expression (Figure 10 A-B).

**Figure 10.**
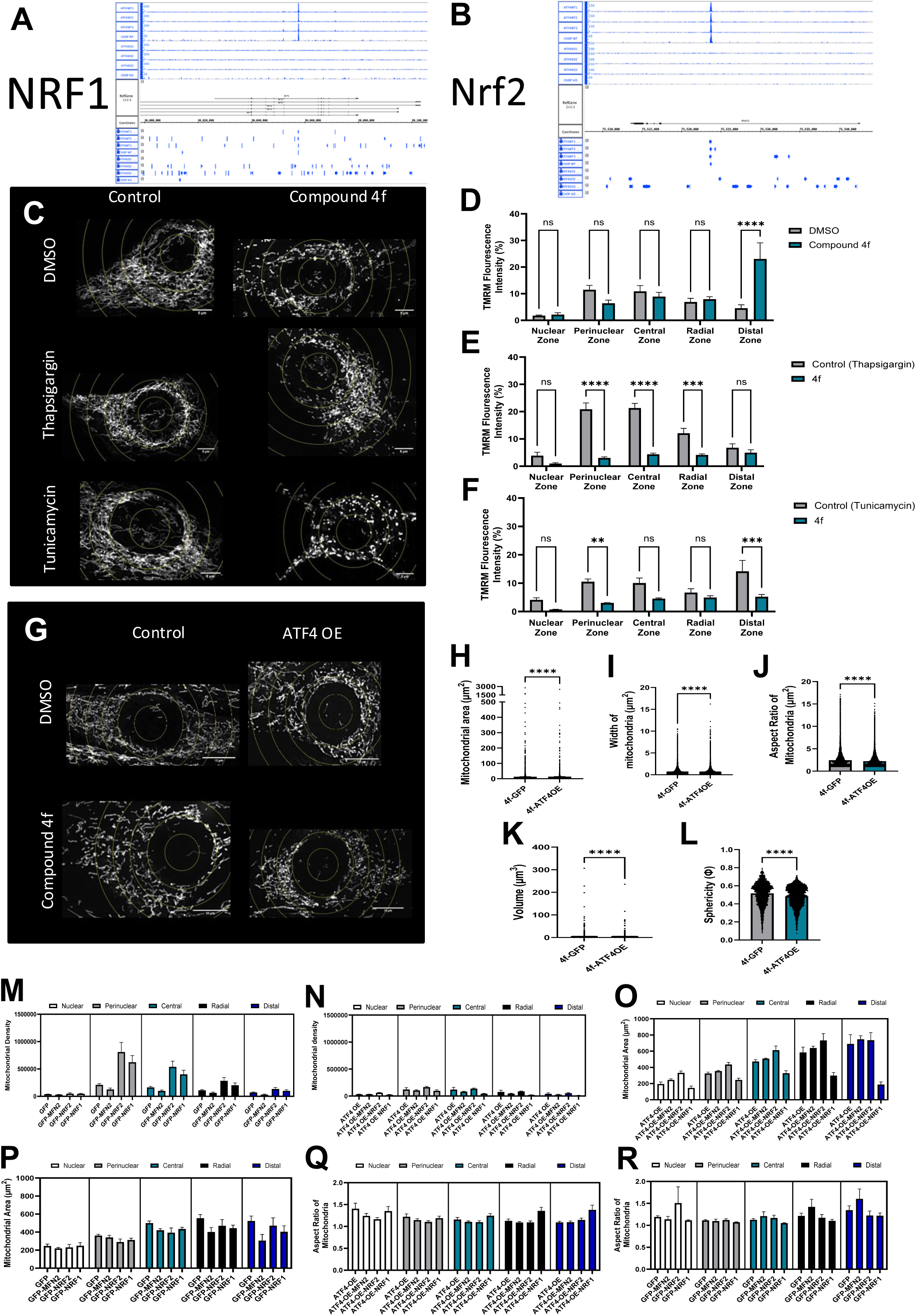
Chromatin immunoprecipitation sequencing (ChIP-Seq) analysis from WT and ATF4 or CHOP deficient MEFs treated with tunicamycin. ATF4 and CHOP bind to promotors of the genes encoding NRF1 (A) and Nrf2 (B) during cellular stress. (C) Representative confocal microscopy images showing mitochondrial distribution (MitoView650, red) and nuclei (DAPI, blue) for C2C12 myoblasts under basal conditions or treated with “Compound 4f,” a small molecule inhibitor of Nrf2 (D-F). Representative confocal images showing mitochondrial morphology (MitoView650, red) in myoblasts from controls and ATF4-overexpressing (ATF4 OE) cultures treated with either DMSO (vehicle) or Compound 4f (G). Analysis of mitochondrial properties in controls compared with ATF4 OE cells under pharmacological treatment (H–L): mitochondrial area (H), mitochondrial width (I), mitochondrial aspect ratio (J), mitochondrial volume (K), mitochondrial sphericity (L). The red bar represents increased mitochondrial properties compared with controls. C2C12 myoblasts were infected with adenovirus expressing GFP or ATF4 with and without the presence of inhibitors of NRF2, Nrf2, or the combination of both. (M–R). Mitochondrial density (arbitrary units) was analyzed as the integrated fluorescence intensity normalized to myotube area in GFP control, NRF1, and NRF2-overexpressing cells (M) and in ATF4 OE cells (N). Mitochondrial area (μm2) was measured as the mean area of mitochondrial structures in GFP, NRF1, NRF2 cells (O) and ATF4 OE cells (P). Mitochondrial aspect ration (μ2) was measured in GFP, NRF1, NRF2 cells (Q) and ATF4 OE cells (R).

#### The small molecule inhibitor of Nrf2, Compound 4f, prevents the formation of Megamitochondria typically seen with induced cellular stress

To determine if Nrf2 is required for the ATF4-dependent, MFN2-mediated mitochondrial changes with induced stress, we treated C2C12 myoblasts with either DMSO, thapsigargin, or tunicamycin, with or without the addition of Compound 4f (Figure 10C). We found that, with the addition of 4f, the mitochondrial hyperpolarization typically observed under induced cellular stress is lost (Figure 10 D-F), indicating a severe disruption of stress-induced mitochondrial adaptations.

#### Overexpression of ATF4 with treatment with 4f shows similar results in the loss of Megamitochondria formation seen with induced cellular stress

To determine whether our results observed with induced cellular stress and treatment with 4f could be recapitulated by overexpressing ATF4, we performed identical experiments as in our induced cellular stress model, except that, instead of chemically inducing cellular stress, we used adenovirus expressing either GFP or ATF4 (Figure 10G). As anticipated, the overexpression of ATF4 showed similar changes in mitochondrial morphology and the loss of Megamitochondria formation observed with induced cellular stress (Figure 10G).

Further quantitative analyses conducted under various experimental settings showed that NRF1 and Nrf2 chemical modulation had varied effects on the size, shape, volume, and mass of mitochondria in GFP and ATF4 OE cells (Figure 10 M-R). Specifically, ATF4-driven alterations in mitochondrial size and complexity were downregulated when the NRF pathway was inhibited, suggesting that ATF4 and NRF signaling interact functionally to control mitochondrial structure. Results from Figure 10 show that the NRF1 and Nrf2 signaling pathways play a crucial role in the mitochondrial remodeling induced by ATF4. These results provide support for the idea that ATF4 coordinates the redistribution of mitochondrial spatial coordinates and ultrastructural remodeling in response to cellular stress by activating transcriptional programs dependent on NRF1 and Nrf2 signaling.

## Discussion

This study sought to investigate the molecular mechanisms behind mitochondrial and MERC changes observed during cellular stress. We found that ATF4 activation is linked to significant alterations in central carbon metabolism in the *Drosophila* model, including coordinated modifications in tricarboxylic acid cycle intermediates such citrate, fumarate, succinate, and malate. Elevation of these metabolites is consistent with higher anaplerotic flux and mitochondrial metabolic throughput, suggesting that mega-mitochondrial formation and mitochondrial expansion are supported by improved metabolic capacity rather than by bioenergetic collapse. This metabolic state is ideal for maintaining larger mitochondrial volumes, increased cristae surface area, and long-range mitochondrial communication via nanotunneling (Vincent, Turnbull et al. 2017, Palese, Rakotobe et al. 2025).

Importantly, ATF4 activation causes mitochondrial expansion as well as significant modification of cristae architecture. Quantitative ultrastructural analysis reveals that increases in mitochondrial size and volume are accompanied by coordinated changes in cristae number, density, and surface area. These findings show that mega-mitochondria remain bioenergetically structured and functionally competent. As a result, cristae remodeling is a functionally integrated adaptation that supports both the energy requirements of huge mitochondrial networks and elevated oxidative metabolism (Ježek, Jabůrek et al. 2023, Kawano, Bazila et al. 2023).

Fly metabolomics also reveals changes in amino acid pools, including branched-chain amino acids, methionine, cysteine, and taurine-related metabolites, which directly impact ISR and redox regulatory pathways. These modifications are believed to under stress. Cysteine- and glutathione-related metabolites are particularly significant in connecting ATF4 activation with NRF2-driven antioxidant pathways (Wallis, Morehead et al. 2021). This supports the idea that regulated redox signaling facilitates mitochondrial expansion, cristae remodeling, and fusion. Conversely, excessive oxidative stress would impede these processes.

The fly model also affects the MERCS by altering several metabolite types. This metabolic rewiring, driven by ATF4, affects the lipid composition, curvature, and fluidity of the membrane, as reflected by changes in glycerolipid- and phospholipid-related intermediates. As a result of these alterations, larger MERCS should become more stable, allowing for more consistent lipid transfer, coordination of mitochondrial fusion machinery, and prolonged calcium exchange (Szymański, Janikiewicz et al. 2017, Bassot, Chen et al. 2021). Consequently, larger MERCS provide a biochemical and functional platform that can enable mega-mitochondrial development, cristae alteration, and the generation of Mitochondrial Nanotunnels (Vincent, Turnbull et al. 2017).

Metabolomic analysis shows ATF4 activation in mammalian myotubes. In our in vitro studies, myotubes show alterations in glycolytic intermediates, pyruvate handling, and TCA cycle flux, suggesting metabolically flexible and oxidatively active state (Philp, Perez-Schindler et al. 2010). These changes are consistent with elevated mitochondrial volume and area, modified cristae structure, and enhanced mitochondrial connectivity, indicating that metabolic remodeling actively enlarges mitochondria (Kawano, Bazila et al. 2023).

Metabolic–calcium–redox pathway seems to be very important for controlling mitochondrial nanotunnel formation (Lavorato, Iyer et al. 2017, Vincent, Turnbull et al. 2017). Requirements for the formation of nanotunnels include remodeling the membrane, coordinate the cytoskeleton, and provide constant energy input (Vincent, Turnbull et al. 2017, Lavorato 2018). The metabolic patterns observed in the fly and myotube models are consistent with the biosynthetic and energetic needs fundamental to the formation and maintenance of nanotunnels. As a potential alternative to stress-induced damage, nanotunneling could be an adaptive, ISR-driven method for transporting metabolites, calcium, and redox signals across widely distributed mitochondria (Vincent, Turnbull et al. 2017).

ATF4 activation reshaped mitochondrial spatial organization. Quantitative spatial analysis revealed redistribution across perinuclear, nuclear-adjacent, radial, and distal cytoplasmic zones. ATF4 overexpression promoted mitochondrial enrichment in perinuclear and radial regions, whereas ATF4 knockdown produced fragmented mitochondria confined to the cell periphery. Perinuclear localization promotes sustained interactions with the endoplasmic reticulum and calcium exchange, whereas radial organization along cytoskeletal tracks enables nanotunnel formation that links distant mitochondrial domains (Vincent, Turnbull et al. 2017). This coordinated zonal positioning is therefore essential for long-range mitochondrial communication and network cohesion.

MFN2 controls fusion and MERC tethering, which connects changes in cristae shape, mitochondrial size, and nanotunneling (Vincent, Turnbull et al. 2017, Zaman and Shutt 2022). These structural changes are interconnected, as MFN2 disruption abolishes mitochondrial extension, nanotunneling, and zonal redistribution (Sood 2015, Vincent, Turnbull et al. 2017). Calcium and redox signaling appear to function within this framework to couple metabolic output to membrane dynamics and cytoskeletal interactions required for large-scale mitochondrial remodeling. Comparison of in vivo and in vitro systems highlight how physiological context modulates ATF4-dependent regulation: systemic inputs in vivo broaden metabolic engagement downstream of ATF4, strengthening lipid and redox pathway involvement and promoting sustained mega-mitochondrial formation across tissues. Therefore, ATF4-driven ISR signaling is sufficient to alter mitochondrial size, cristae architecture, spatial organization and connectivity with activation.

Collectively, ATF4 is established as a master regulator that integrates ISR and metabolic state, calcium and redox signaling, and spatial organization to control mitochondrial architecture at several scales. ATF4 establishes ISR regulation as a key factor in determining mitochondrial health in physiology and disease by controlling MERC dynamics and metabolic flux, which determines whether Megamitochondria and mitochondrial nanotunneling serve adaptive stress responses or become maladaptive under chronic or pathological conditions.

## Data availability statement

All data generated or analyzed during this study are included in this published article and its Supplementary information files. Additional data can be requested from the corresponding author.

## Acknowledgments

UNCF/Bristol-Myers Squibb E.E. Just Faculty Fund, Career Award at the Scientific Interface (CASI Award) from the Burroughs Welcome Fund (BWF) ID # 1021868.01, BWF Ad-hoc Award, NIH Small Research Pilot Subaward 5R25HL106365-12 from the National Institutes of Health PRIDE Program, DK020593, Vanderbilt Diabetes and Research Training Center for DRTC Alzheimer’s Disease Pilot & Feasibility Program, CZI Science Diversity Leadership grant number 2022-253529 from the Chan Zuckerberg Initiative DAF, an advised fund of the Silicon Valley Community Foundation to A.H.J. NSF NRT grant 19-22697 (to A.C.), NSF BPE grant 22-17621 (to A.C.), and the Asness Family Fund through the Frist Center for Autism & Innovation (K. Stassun, PI) (to A.C.) This work was also supported by National Institutes of Health HL006221 (BG).

## Supplementary File

**Supplementary Figure 1.** This genome browser view represents ChIP-seq signal tracks across the CHOP and BIP gene locuses under ATF4 KO and ATFE control conditions. The blue peaks indicate regions of enriched DNA binding from immunoprecipitation.

**Supplementary Video 1.** Control indirect flight muscle mitochondrial network (low magnification, 10 µm). 3D reconstruction of Drosophila indirect flight muscle fibers showing the full mitochondrial network. Muscle fibers are shown in red and mitochondria in green, with segmentation of all mitochondria within the imaged volume.

**Supplementary Video 2.** ATF4 knockout indirect flight muscle mitochondrial network (low magnification, 10 µm). 3D reconstruction of Drosophila indirect flight muscle fibers lacking ATF4 showing the full mitochondrial network. Muscle fibers are shown in red and mitochondria in green, with segmentation of all mitochondria within the imaged volume.

**Supplementary Video 3.** Control indirect flight muscle mitochondrial network (intermediate magnification, 7 µm). Higher magnification 3D reconstruction of Drosophila indirect flight muscle fibers illustrating mitochondrial organization between fibers. Muscle fibers are shown in red and mitochondria in green, with segmentation of all mitochondria within the imaged volume.

**Supplementary Video 4.** ATF4 knockout indirect flight muscle mitochondrial network (intermediate magnification, 7 µm). Higher magnification 3D reconstruction of ATF4 knockout Drosophila indirect flight muscle fibers illustrating mitochondrial organization. Muscle fibers are shown in red and mitochondria in green, with segmentation of all mitochondria within the imaged volume.

**Supplementary Video 5.** Control mitochondrial architecture in indirect flight muscle (high magnification, 7 µm). High-resolution 3D segmentation of the mitochondrial network in Drosophila indirect flight muscle, showing all mitochondria (green) within the fiber volume.

**Supplementary Video 6.** ATF4 knockout mitochondrial architecture in indirect flight muscle (high magnification, 7 µm). High-resolution 3D segmentation of the mitochondrial network in ATF4 knockout Drosophila indirect flight muscle, showing all mitochondria (green) within the fiber volume.

**Supplementary Video 7.** Control indirect flight muscle mitochondrial network (low magnification, 10 µm). 3D reconstruction of Drosophila indirect flight muscle fibers showing the organization of mitochondria between contractile fibers. Muscle fibers are shown in red and mitochondria in green, with segmentation of all mitochondria within the imaged volume.

**Supplementary Video 8.** ATF4 overexpression indirect flight muscle mitochondrial network (low magnification, 10 µm). 3D reconstruction of Drosophila indirect flight muscle fibers with ATF4 overexpression (ATF4 OE) showing mitochondrial organization between fibers. Muscle fibers are shown in red and mitochondria in green, with segmentation of all mitochondria within the imaged volume.

**Supplementary Video 9.** Control indirect flight muscle mitochondrial architecture (intermediate magnification, 10 µm). Higher-magnification 3D reconstruction of control Drosophila indirect flight muscle, illustrating mitochondrial alignment along muscle fibers. Muscle fibers are shown in red and mitochondria in green, with segmentation of all mitochondria within the imaged volume.

**Supplementary Video 10.** ATF4 overexpression indirect flight muscle mitochondrial architecture (intermediate magnification, 10 µm). Higher-magnification 3D reconstruction of ATF4 OE Drosophila indirect flight muscle, illustrating mitochondrial alignment and network organization. Muscle fibers are shown in red and mitochondria in green, with segmentation of all mitochondria within the imaged volume.

**Supplementary Video 11.** Control mitochondrial network segmentation in indirect flight muscle (mitochondria only, 10 µm). 3D segmentation showing all mitochondria (green) within control Drosophila indirect flight muscle fibers, highlighting the mitochondrial network architecture throughout the reconstructed volume.

**Supplementary Video 12.** ATF4 overexpression mitochondrial network segmentation in indirect flight muscle (mitochondria only, 10 µm). 3D segmentation showing all mitochondria (green) within ATF4 OE Drosophila indirect flight muscle fibers, illustrating the mitochondrial network architecture within the reconstructed volume.

**Supplementary Video 13** - Single mitochondrion reconstruction from wild-type *Drosophila* indirect flight muscle, showing a compact mitochondrial morphology with short nanotunnel connections.

**Supplementary Video 14** - Single mitochondrion reconstruction from ATF4 knockout (KO) *Drosophila* indirect flight muscle, showing elongated mitochondrial morphology with extended nanotunnels connecting distal mitochondrial regions.

**Supplementary Video 15** - Single mitochondrion reconstruction from ATF4 overexpression (OE) *Drosophila* indirect flight muscle, displaying enlarged and clustered mitochondrial structures with nanotunnel-like protrusions.

## References

Adams, C. M., S. M. Ebert and M. C. Dyle (2017). "Role of ATF4 in skeletal muscle atrophy." Curr Opin Clin Nutr Metab Care 20(3): 164–168.

Bartsakoulia, M., A. Pyle, D. Troncoso-Chandía, J. Vial-Brizzi, M. V. Paz-Fiblas, J. Duff, H. Griffin, V. Boczonadi, H. Lochmüller, S. Kleinle, P. F. Chinnery, S. Grünert, J. Kirschner, V. Eisner and R. Horvath (2018). "A novel mechanism causing imbalance of mitochondrial fusion and fission in human myopathies." Hum Mol Genet 27(7): 1186–1195.

Bassot, A., J. Chen and T. Simmen (2021). "Post-translational modification of cysteines: a key determinant of endoplasmic reticulum-mitochondria contacts (MERCs)." Contact 4: 25152564211001213.

Boudina, S., S. Sena, H. Theobald, X. Sheng, J. J. Wright, X. X. Hu, S. Aziz, J. I. Johnson, H. Bugger, V. G. Zaha and E. D. Abel (2007). "Mitochondrial energetics in the heart in obesity-related diabetes: direct evidence for increased uncoupled respiration and activation of uncoupling proteins." Diabetes 56(10): 2457–2466.

Bratic, A. and N. G. Larsson (2013). "The role of mitochondria in aging." J Clin Invest 123(3): 951–957.

Bustos, G., P. Cruz, A. Lovy and C. Cárdenas (2017). "Endoplasmic Reticulum-Mitochondria Calcium Communication and the Regulation of Mitochondrial Metabolism in Cancer: A Novel Potential Target." Front Oncol 7: 199.

Chan, D. C. (2012). "Fusion and fission: interlinked processes critical for mitochondrial health." Annu Rev Genet 46: 265–287.

Cogliati, S., J. A. Enriquez and L. Scorrano (2016). "Mitochondrial Cristae: Where Beauty Meets Functionality." Trends Biochem Sci 41(3): 261–273.

Del Campo, A., I. Contreras-Hernández, M. Castro-Sepúlveda, C. A. Campos, R. Figueroa, M. F. Tevy, V. Eisner, M. Casas and E. Jaimovich (2018). "Muscle function decline and mitochondria changes in middle age precede sarcopenia in mice." Aging (Albany NY) 10(1): 34–55.

Dong, Z. and X. Yao (2022). "Insight of the role of mitochondrial calcium homeostasis in hepatic insulin resistance." Mitochondrion 62: 128–138.

Ebert, S. M., M. C. Dyle, S. D. Kunkel, S. A. Bullard, K. S. Bongers, D. K. Fox, J. M. Dierdorff, E. D. Foster and C. M. Adams (2012). "Stress-induced skeletal muscle Gadd45a expression reprograms myonuclei and causes muscle atrophy." J Biol Chem 287(33): 27290–27301.

Garza-Lopez, E., Z. Vue, P. Katti, K. Neikirk, M. Biete, J. Lam, H. K. Beasley, A. G. Marshall, T. A. Rodman, T. A. Christensen, J. L. Salisbury, L. Vang, M. Mungai, S. AshShareef, S. A. Murray, J. Shao, J. Streeter, B. Glancy, R. O. Pereira, E. D. Abel and A. Hinton, Jr. (2021). "Protocols for Generating Surfaces and Measuring 3D Organelle Morphology Using Amira." Cells 11(1).

Glancy, B. (2020). "Visualizing Mitochondrial Form and Function within the Cell." Trends Mol Med 26(1): 58–70.

Hall, H., P. Medina, D. A. Cooper, S. E. Escobedo, J. Rounds, K. J. Brennan, C. Vincent, P. Miura, R. Doerge and V. M. Weake (2017). "Transcriptome profiling of aging Drosophila photoreceptors reveals gene expression trends that correlate with visual senescence." BMC Genomics 18(1): 894.

Han, J., S. H. Back, J. Hur, Y. H. Lin, R. Gildersleeve, J. Shan, C. L. Yuan, D. Krokowski, S. Wang, M. Hatzoglou, M. S. Kilberg, M. A. Sartor and R. J. Kaufman (2013). "ER-stress-induced transcriptional regulation increases protein synthesis leading to cell death." Nat Cell Biol 15(5): 481–490.

Hara, Y., F. Yuk, R. Puri, W. G. Janssen, P. R. Rapp and J. H. Morrison (2014). "Presynaptic mitochondrial morphology in monkey prefrontal cortex correlates with working memory and is improved with estrogen treatment." Proc Natl Acad Sci U S A 111(1): 486–491.

Hinton, A., Jr., S. M. Claypool, K. Neikirk, N. Senoo, C. N. Wanjalla, A. Kirabo and C. R. Williams (2024). "Mitochondrial Structure and Function in Human Heart Failure." Circ Res 135(2): 372–396.

Hinton, A., Jr., P. Katti, T. A. Christensen, M. Mungai, J. Shao, L. Zhang, S. Trushin, A. Alghanem, A. Jaspersen, R. E. Geroux, K. Neikirk, M. Biete, E. G. Lopez, B. Shao, Z. Vue, L. Vang, H. K. Beasley, A. G. Marshall, D. Stephens, S. Damo, J. Ponce, C. K. E. Bleck, I. Hicsasmaz, S. A. Murray, R. A. C. Edmonds, A. Dajles, Y. D. Koo, S. Bacevac, J. L. Salisbury, R. O. Pereira, B. Glancy, E. Trushina and E. D. Abel (2023). "A Comprehensive Approach to Sample Preparation for Electron Microscopy and the Assessment of Mitochondrial Morphology in Tissue and Cultured Cells." Adv Biol (Weinh) 7(10): e2200202.

Hinton, A., Jr., P. Katti, M. Mungai, D. D. Hall, O. Koval, J. Shao, Z. Vue, E. G. Lopez, R. Rostami, K. Neikirk, J. Ponce, J. Streeter, B. Schickling, S. Bacevac, C. Grueter, A. Marshall, H. K. Beasley, Y. Do Koo, S. C. Bodine, N. G. R. Nava, A. M. Quintana, L. S. Song, I. M. Grumbach, R. O. Pereira, B. Glancy and E. D. Abel (2024). "ATF4-dependent increase in mitochondrial-endoplasmic reticulum tethering following OPA1 deletion in skeletal muscle." J Cell Physiol 239(4): e31204.

Jenkins, B. C., K. Neikirk, P. Katti, S. M. Claypool, A. Kirabo, M. R. McReynolds and A. Hinton, Jr. (2024). "Mitochondria in disease: changes in shapes and dynamics." Trends Biochem Sci 49(4): 346–360.

Ježek, P., M. Jabůrek, B. Holendová, H. Engstová and A. Dlasková (2023). "Mitochondrial cristae morphology reflecting metabolism, superoxide formation, redox homeostasis, and pathology." Antioxidants & Redox Signaling 39(10-12): 635–683.

Katti, P., M. Rai, S. Srivastava, P. D’Silva and U. Nongthomba (2021). "Marf-mediated mitochondrial fusion is imperative for the development and functioning of indirect flight muscles (IFMs) in drosophila." Exp Cell Res 399(2): 112486.

Kawano, I., B. Bazila, P. Ježek and A. Dlasková (2023). "Mitochondrial dynamics and cristae shape changes during metabolic reprogramming." Antioxidants & Redox Signaling 39(10-12): 684–707.

Lam, J., P. Katti, M. Biete, M. Mungai, S. AshShareef, K. Neikirk, E. Garza Lopez, Z. Vue, T. A. Christensen, H. K. Beasley, T. A. Rodman, S. A. Murray, J. L. Salisbury, B. Glancy, J. Shao, R. O. Pereira, E. D. Abel and A. Hinton, Jr. (2021). "A Universal Approach to Analyzing Transmission Electron Microscopy with ImageJ." Cells 10(9).

Lavorato, M. (2018). "Microtubules and mitochondria nanotunnels." Physiological Mini Reviews 11.

Lavorato, M., V. R. Iyer, W. Dewight, R. R. Cupo, V. Debattisti, L. Gomez, S. De la Fuente, Y.-T. Zhao, H. H. Valdivia and G. Hajnóczky (2017). "Increased mitochondrial nanotunneling activity, induced by calcium imbalance, affects intermitochondrial matrix exchanges." Proceedings of the National Academy of Sciences 114(5): E849–E858.

Mustafi, D., S. Kikano and K. Palczewski (2014). "Serial block face-scanning electron microscopy: a method to study retinal degenerative phenotypes." Curr Protoc Mouse Biol 4(4): 197–204.

Neikirk, K., E. G. Lopez, A. G. Marshall, A. Alghanem, E. Krystofiak, B. Kula, N. Smith, J. Shao, P. Katti and A. Hinton, Jr. (2023). "Call to action to properly utilize electron microscopy to measure organelles to monitor disease." Eur J Cell Biol 102(4): 151365.

Pakos-Zebrucka, K., I. Koryga, K. Mnich, M. Ljujic, A. Samali and A. M. Gorman (2016). "The integrated stress response." EMBO Rep 17(10): 1374–1395.

Palese, F., M. Rakotobe and C. Zurzolo (2025). "Transforming the concept of connectivity: unveiling tunneling nanotube biology and their roles in brain development and neurodegeneration." Physiological Reviews 105(3): 1823–1865.

Patra, A. K., Y. M. Kwon, S. G. Kang, Y. Fujiwara and S. J. Kim (2016). "The complete mitochondrial genome sequence of the tubeworm Lamellibrachia satsuma and structural conservation in the mitochondrial genome control regions of Order Sabellida." Mar Genomics 26: 63–71.

Pereira, R. O., S. M. Tadinada, F. M. Zasadny, K. J. Oliveira, K. M. P. Pires, A. Olvera, J. Jeffers, R. Souvenir, R. McGlauflin, A. Seei, T. Funari, H. Sesaki, M. J. Potthoff, C. M. Adams, E. J. Anderson and E. D. Abel (2017). "OPA1 deficiency promotes secretion of FGF21 from muscle that prevents obesity and insulin resistance." Embo j 36(14): 2126–2145.

Philp, A., J. Perez-Schindler, C. Green, D. L. Hamilton and K. Baar (2010). "Pyruvate suppresses PGC1α expression and substrate utilization despite increased respiratory chain content in C2C12 myotubes." American Journal of Physiology-Cell Physiology 299(2): C240–C250.

Rasmussen, B. B. and C. M. Adams (2020). "ATF4 is a fundamental regulator of nutrient sensing and protein turnover." The Journal of Nutrition 150(5): 979–980.

Schneider, C. A., W. S. Rasband and K. W. Eliceiri (2012). "NIH Image to ImageJ: 25 years of image analysis." Nat Methods 9(7): 671–675.

Sood, A. (2015). "Wiring the adaptive response of mitochondria to metabolic transitions: a Mitofusin-2-dependent proteolytic elimination of OPA1 accompanies cristae and mitochondria-ER contacts remodelling in the postprandial mouse liver."

Szymański, J., J. Janikiewicz, B. Michalska, P. Patalas-Krawczyk, M. Perrone, W. Ziółkowski, J. Duszyński, P. Pinton, A. Dobrzyń and M. R. Więckowski (2017). "Interaction of mitochondria with the endoplasmic reticulum and plasma membrane in calcium homeostasis, lipid trafficking and mitochondrial structure." International Journal of Molecular Sciences 18(7): 1576.

Tameire, F., Verginadis, II, N. M. Leli, C. Polte, C. S. Conn, R. Ojha, C. Salas Salinas, F. Chinga, A. M. Monroy, W. Fu, P. Wang, A. Kossenkov, J. Ye, R. K. Amaravadi, Z. Ignatova, S. Y. Fuchs, J. A. Diehl, D. Ruggero and C. Koumenis (2019). "ATF4 couples MYC-dependent translational activity to bioenergetic demands during tumour progression." Nat Cell Biol 21(7): 889–899.

Vincent, A. E., D. M. Turnbull, V. Eisner, G. Hajnóczky and M. Picard (2017). "Mitochondrial nanotunnels." Trends in cell biology 27(11): 787–799.

Vue, Z., K. Neikirk, L. Vang, E. Garza-Lopez, T. A. Christensen, J. Shao, J. Lam, H. K. Beasley, A. G. Marshall, A. Crabtree, J. Anudokem, Jr., B. Rodriguez, B. Kirk, S. Bacevac, T. Barongan, B. Shao, D. C. Stephens, K. Kabugi, H. J. Koh, A. Koh, C. S. Evans, B. Taylor, A. K. Reddy, T. Miller-Fleming, K. V. Actkins, E. Zaganjor, N. Daneshgar, S. A. Murray, B. C. Mobley, S. M. Damo, J. A. Gaddy, B. Riggs, C. Wanjalla, A. Kirabo, M. McReynolds, J. A. Gomez, M. A. Phillips, V. Exil, D. F. Dai and A. Hinton, Jr. (2023). "Three-dimensional mitochondria reconstructions of murine cardiac muscle changes in size across aging." Am J Physiol Heart Circ Physiol 325(5): H965–h982.

Wallis, K. F., L. C. Morehead, J. T. Bird, S. D. Byrum and I. R. Miousse (2021). "Differences in cell death in methionine versus cysteine depletion." Environmental and molecular mutagenesis 62(3): 216–226.

Wende, A. R., B. T. O’Neill, H. Bugger, C. Riehle, J. Tuinei, J. Buchanan, K. Tsushima, L. Wang, P. Caro, A. Guo, C. Sloan, B. J. Kim, X. Wang, R. O. Pereira, M. A. McCrory, B. G. Nye, G. A. Benavides, V. M. Darley-Usmar, T. Shioi, B. C. Weimer and E. D. Abel (2015). "Enhanced cardiac Akt/protein kinase B signaling contributes to pathological cardiac hypertrophy in part by impairing mitochondrial function via transcriptional repression of mitochondrion-targeted nuclear genes." Mol Cell Biol 35(5): 831–846.

Zaman, M. and T. E. Shutt (2022). "The role of impaired mitochondrial dynamics in MFN2-mediated pathology." Frontiers in cell and developmental biology 10: 858286.

Zou, Z., T. Ohta, F. Miura and S. Oki (2022). "ChIP-Atlas 2021 update: a data-mining suite for exploring epigenomic landscapes by fully integrating ChIP-seq, ATAC-seq and Bisulfite-seq data." Nucleic Acids Res 50(W1): W175–w182.

