## Supplementary figures and images for "Stress-Encoded Mitochondrial Plasticity: ATF4 Control of Mega-Mitochondria and Nanotunnel Communication"

### Supplemetary File

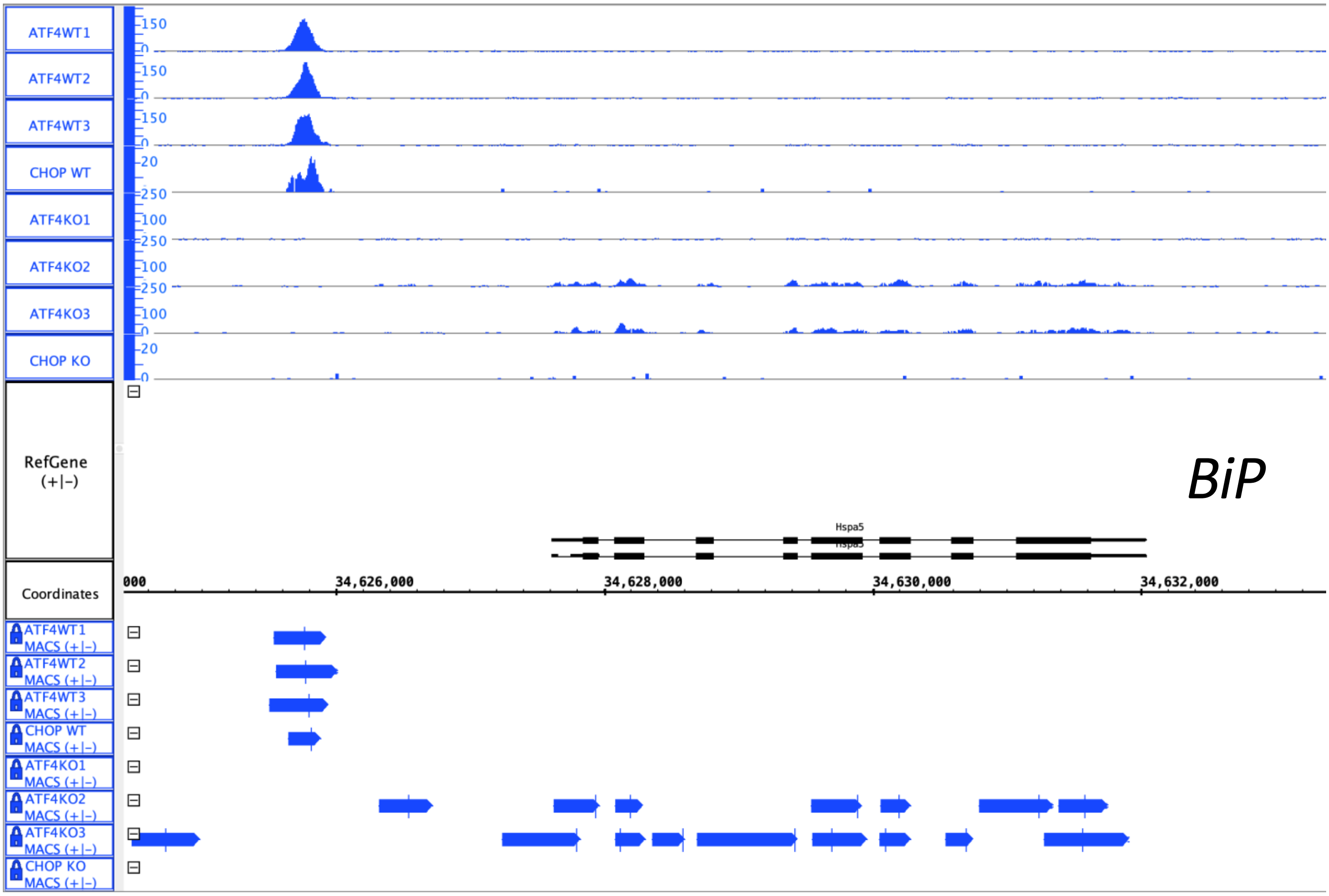

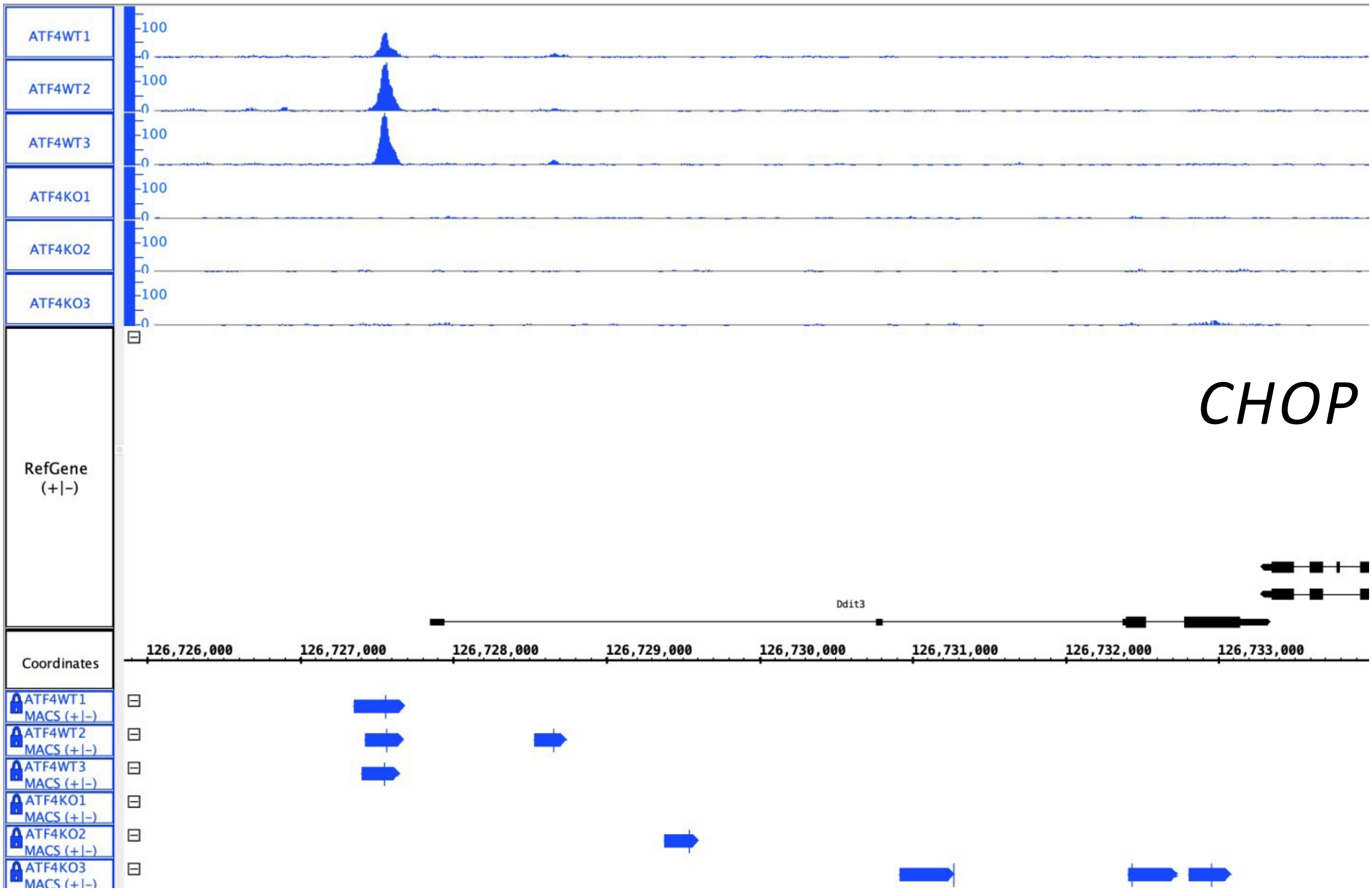

WT

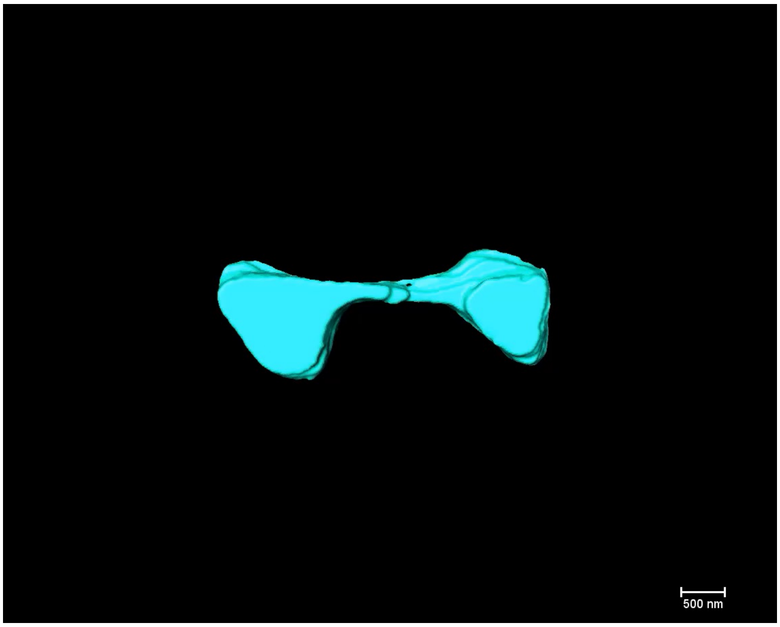


OE

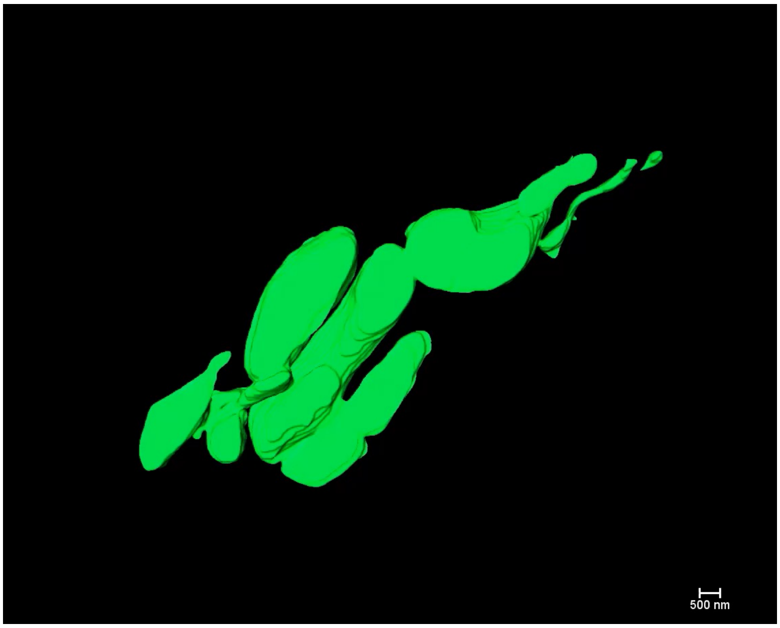


KO

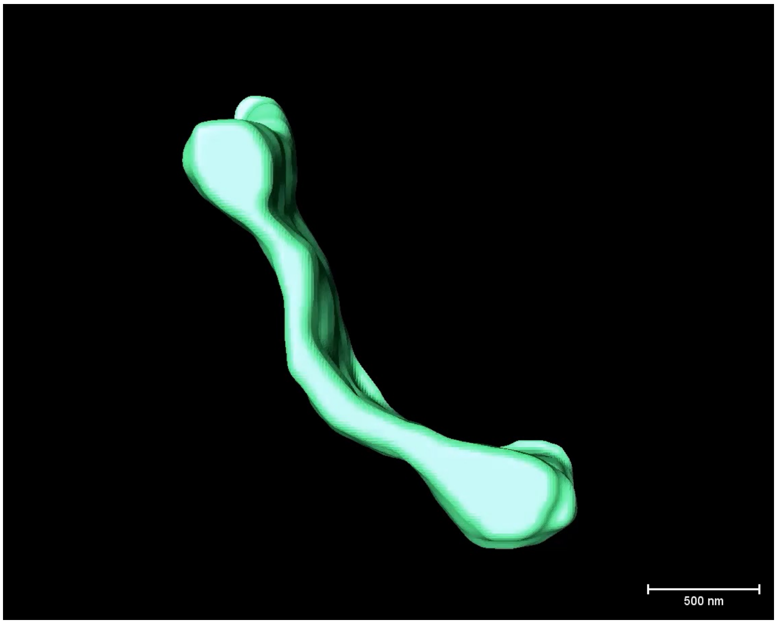

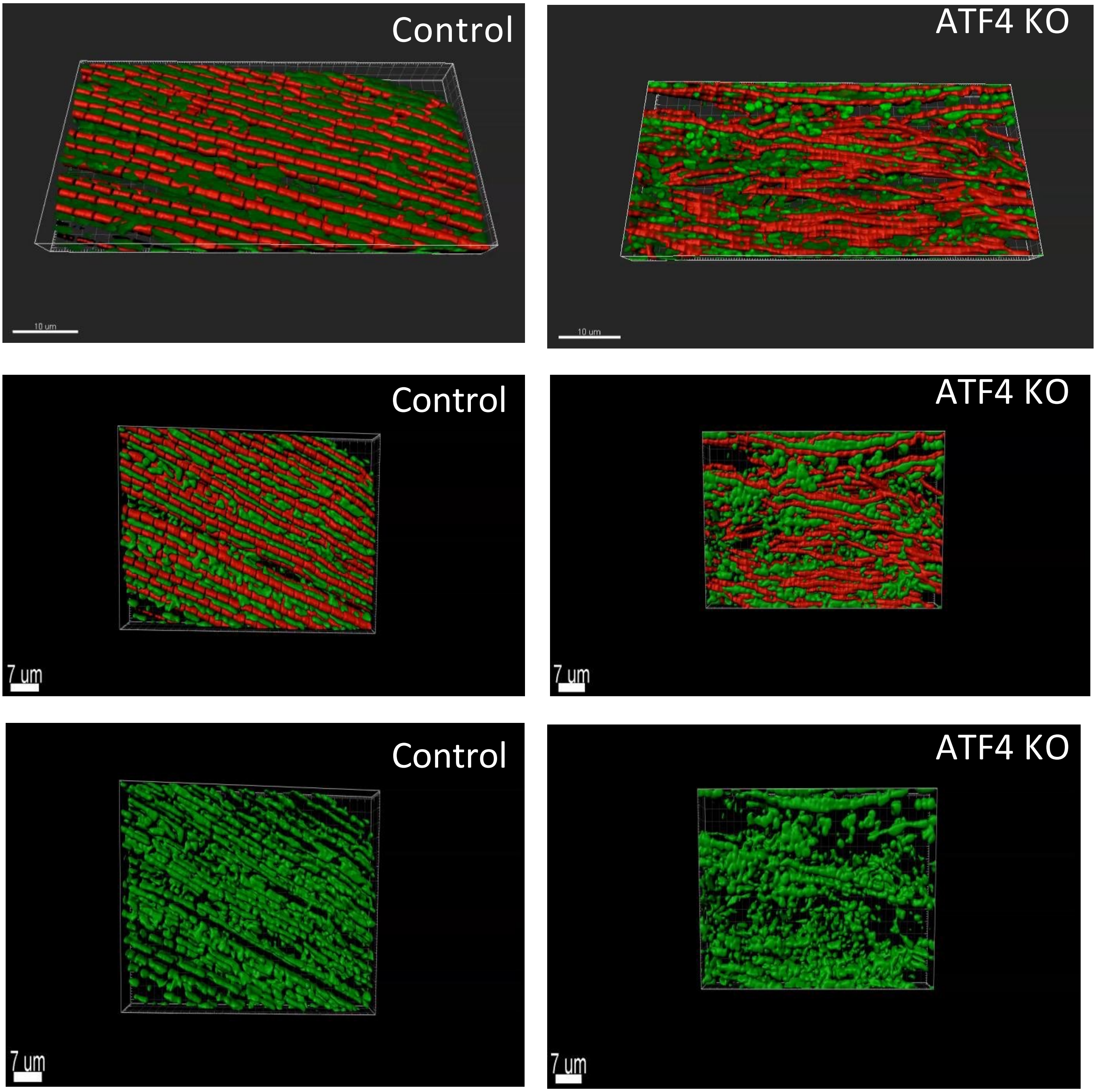

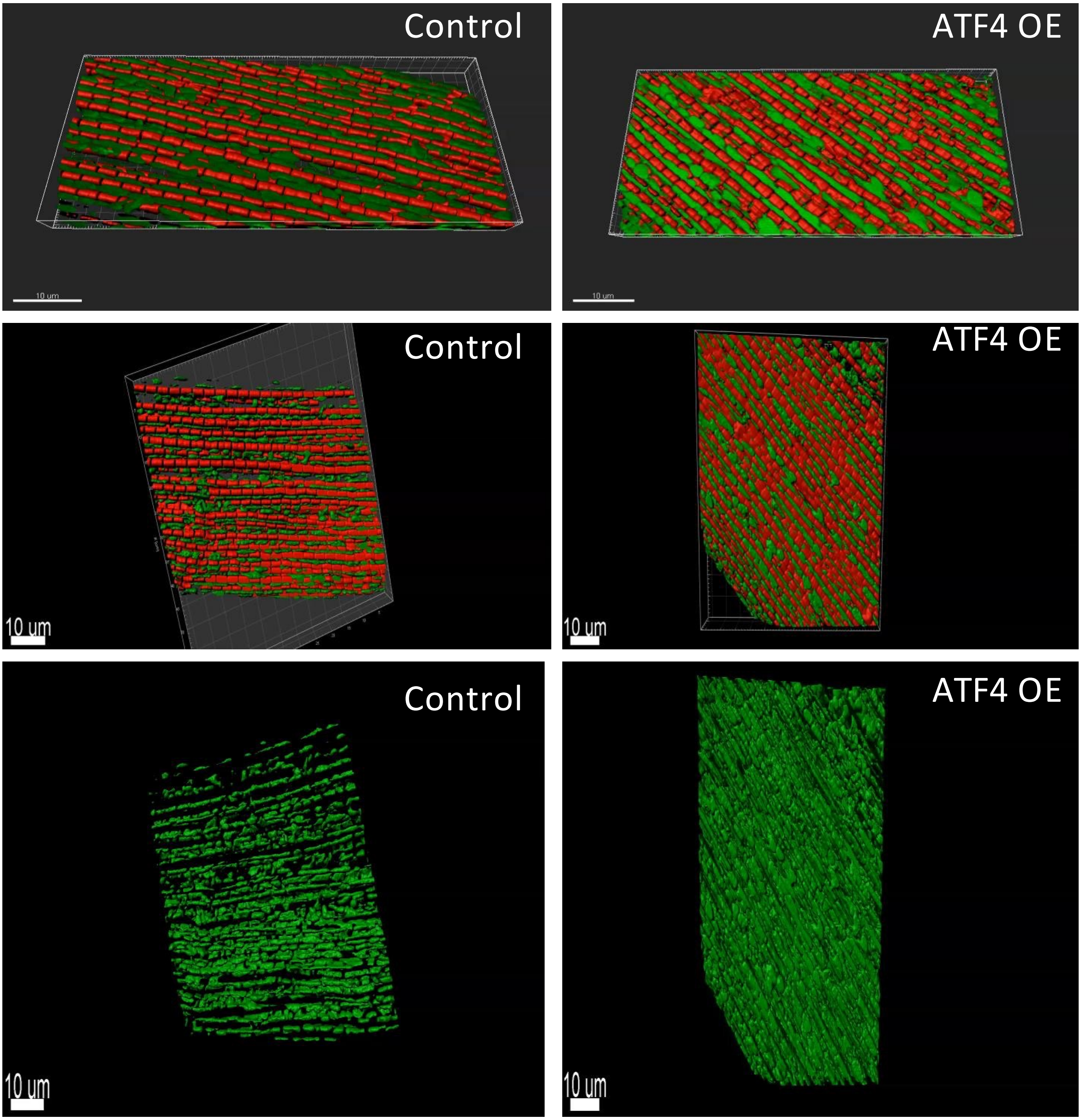
